# Naturalistic Object Representations Depend on Distance and Size Cues

**DOI:** 10.1101/2024.03.16.585308

**Authors:** Grant T. Fairchild, Desiree E. Holler, Sara Fabbri, Michael A. Gomez, Jacqueline C. Walsh-Snow

## Abstract

Egocentric distance and real-world size are important cues for object perception and action. Nevertheless, most studies of human vision rely on two-dimensional pictorial stimuli that convey ambiguous distance and size information. Here, we use fMRI to test whether pictures are represented differently in the human brain from real, tangible objects that convey unambiguous distance and size cues. Participants directly viewed stimuli in two display formats (real objects and matched printed pictures of those objects) presented at different egocentric distances (near and far). We measured the effects of format and distance on fMRI response amplitudes and response patterns. We found that fMRI response amplitudes in the lateral occipital and posterior parietal cortices were stronger overall for real objects than for pictures. In these areas and many others, including regions involved in action guidance, responses to real objects were stronger for near vs. far stimuli, whereas distance had little effect on responses to pictures—suggesting that distance determines relevance to action for real objects, but not for pictures. Although stimulus distance especially influenced response patterns in dorsal areas that operate in the service of visually guided action, distance also modulated representations in ventral cortex, where object responses are thought to remain invariant across contextual changes. We observed object size representations for both stimulus formats in ventral cortex but predominantly only for real objects in dorsal cortex. Together, these results demonstrate that whether brain responses reflect physical object characteristics depends on whether the experimental stimuli convey unambiguous information about those characteristics.

**Significance Statement:** Classic frameworks of vision attribute perception of inherent object characteristics, such as size, to the ventral visual pathway, and processing of spatial characteristics relevant to action, such as distance, to the dorsal visual pathway. However, these frameworks are based on studies that used projected images of objects whose actual size and distance from the observer were ambiguous. Here, we find that when object size and distance information in the stimulus is less ambiguous, these characteristics are widely represented in both visual pathways. Our results provide valuable new insights into the brain representations of objects and their various physical attributes in the context of naturalistic vision.

## Introduction

The neural mechanisms that support object perception in humans have been studied primarily using two-dimensional, pictorial stimuli. While such research has provided valuable insights into human vision, its ecological validity depends on whether pictures elicit similar responses as real, tangible objects perceived in a three-dimensional environment (Snow and Culham, 2021). Multipronged evidence from studies of behavior suggests that real objects are processed differently than pictures. For example, compared to 2-D pictures, real objects facilitate perception (Romero et al., 2018; Holler et al., 2020) and recognition (Ratcliff and Newcombe, 1982; Riddoch and Humphreys, 1987; Chainay and Humphreys, 2001; Farah, 2004; Turnbull et al., 2004; Holler et al., 2019), attract attention (Mustafar et al., 2015; Gerhard et al., 2016; Gomez et al., 2018; Korisky et al., 2019; Korisky and Mudrik, 2021; Sensoy et al., 2021), enhance memory (Snow et al., 2014, 2023), increase valuation (Bushong et al., 2010; Romero et al., 2018), and modulate executive function (Beaucage et al., 2020) and social cognition (Kingstone, 2009; Pönkänen et al., 2011; Risko et al., 2012; Kuhn et al., 2016).

Importantly, real objects elicit different responses from both 2-D and 3-D stereoscopic pictures (Gomez et al., 2018; Holler et al., 2019, 2020; Snow et al., 2023), indicating that response differences across formats cannot be explained by differences in the availability of stereoscopic depth cues (Snow and Culham, 2021). Nevertheless, depth cues provide a gateway to other critical object characteristics, including physical distance and real-world size. Partly through depth cues, real objects convey definite information about their physical distance from the observer (i.e., egocentric distance), and thus their absolute size is also known. In contrast, although the distance to a picture’s surface is known, the distance of the object depicted within the picture is not known (Proffitt, 2006); thus, its physical size is ambiguous and must be inferred. Importantly, egocentric distance and real-world size determine whether a real object is actable: Only appropriately sized objects positioned within reach can be grasped. Accordingly, behavioral and EEG studies show that responses to real objects are influenced by egocentric distance (Gomez et al., 2018; Snow et al., 2023) and actability (Bushong et al., 2010; Fairchild et al., 2021), as well as real-world size (Holler et al., 2019, 2020; Sensoy et al., 2021), whereas responses to pictures are not. Despite accumulating evidence that display format and stimulus distance combine to influence behavior, none of the neuroimaging studies directly comparing real objects and pictures (Fairchild et al., 2021; Freud et al., 2018; Marini et al., 2019; Snow et al., 2011) have examined the effect of distance on object representations. We might expect to observe greater format-dependent effects of distance in dorsal brain areas that operate in the service of action and spatial representation (Mishkin and Ungerleider, 1982; Goodale and Milner, 1992; Vallar et al., 1999) than in ventral areas, where object representations are thought to remain stable (i.e., ‘invariant’) across changes in viewpoint and size (Grill-Spector et al., 1999; James et al., 2002; Vuilleumier et al., 2002).

We used fMRI to compare the effects of egocentric distance on neural responses to real objects and closely matched pictures by presenting the two display formats either within or beyond reachable space. We utilized a rapid event-related design suitable both for univariate comparisons of response amplitudes, and for multivariate analyses of response patterns. We used univariate analyses to assess the effects of distance, display format, and the interaction between the two on BOLD activation. Next, we used multivariate analyses to characterize the contributions of object size and other stimulus features towards patterns of activation elicited by stimuli in each format, at each distance. We predicted that egocentric distance would have a greater effect for real objects than for pictures on BOLD response amplitudes and activation patterns, especially in dorsal brain regions implicated in spatial vision and the perceptual guidance of action.

## Materials and Methods

**Participants.** 23 healthy adults participated in the experiment (18 females, 5 males; mean age = 25.57 years, SD = 4.43). Three participants’ data were excluded either due to technical issues or because the participant did not complete the scanning session (17 females, 3 males; mean age = 25.10, SD = 4.38). Each participant’s data was collected during a single scanning session. Participants received $50 per hour in compensation. All participants were right-handed, had normal or corrected-to-normal visual acuity, and provided informed consent in accordance with the University of Nevada, Reno Institutional Review Board.

### Stimuli and Apparatus

In a rapid event-related design, participants directly viewed real-world objects and matched high-resolution 2-D printed pictures of the same items presented at two egocentric locations in the scanner (**Figure 1**). The objects were pairs of everyday tools, including scissors, tongs; key, screwdriver; hammer, cleaver; paint roller, pizza cutter, all presented in their typical real-world size (**Figure 2a-b**). Each of the tools had a similar hand/arm action to one other tool in the stimulus set, thus yielding four pairs of two objects with similar action associations (e.g., a key and a screwdriver both use a rotating hand motion, while a paint roller and a pizza cutter both use a back-and-forth arm motion). We used two different exemplars of each item (e.g., two different keys of varied colors and shapes) to limit decoding based on lower-level stimulus attributes, such as color, and to minimize repetition effects within runs (Grill-Spector et al., 2006). In cases where the tools would normally include ferrous components incompatible with the magnetic field of the MR environment, prop or plastic substitutes were used instead. The real objects were mounted on semicircular black foam core boards that were cut to match the upper circumference of the magnet bore. The stimuli were mounted with the handles oriented rightward, towards the participants’ dominant (right) hands. The picture stimuli were high-resolution matte color photographs of each of the sixteen real tool exemplars photographed using a Canon Rebel T2i DSLR camera with constant F-stop and shutter speed. The photographs were taken with the real objects mounted on the viewing platform, with the platform fixed at a comparable angle and viewing distance to participant’s view in the MRI scanner, under lighting conditions matching those in the scanner. The photos were matched closely to their real-world counterparts for brightness, shading, position, orientation, retinal extent, and background (Fairchild et al., 2021) as displayed during the experiment. To ensure that stimulus position and orientation were held constant across trials, we attached a wooden pedestal to the underside of each stimulus plate that fitted into a slot in the viewing platform (described below). The photos were printed and then mounted to foam core boards. Photos were printed to match the size and shape of the real object stimuli. All stimuli and equipment were pilot-tested in the magnet with an inert phantom to ensure that they did not produce artifacts in the MR signal. We conducted analogous tests of our protocol with an inert phantom to ensure that motion during manual placement of the stimuli did not produce artifacts.

**Figure 1:**
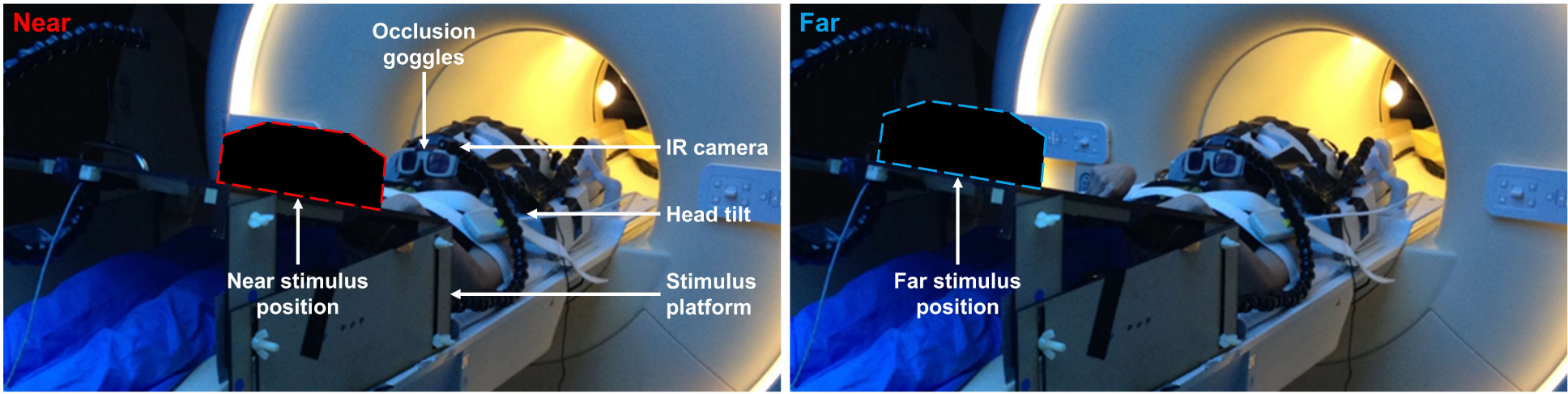
The experimental setup. Participants in the scanner viewed real objects and printed pictures placed on a platform either within reach (left panel) or outside of reach (right panel) of the participants’ hands. The shapes outlined in red and blue show the approximate location and size of stimulus mounts at the near and far positions respectively. Participants’ heads were tilted in the coil to permit direct viewing of the stimuli, without the use of mirrors. Viewing times were controlled using liquid crystal glasses that alternate between transparent and opaque states. LEDs illuminated the stimuli when the glasses were open. An infrared camera recorded runs so that errors (e.g., manual positioning errors or incorrectly selected stimuli) could be detected.

**Figure 2:**
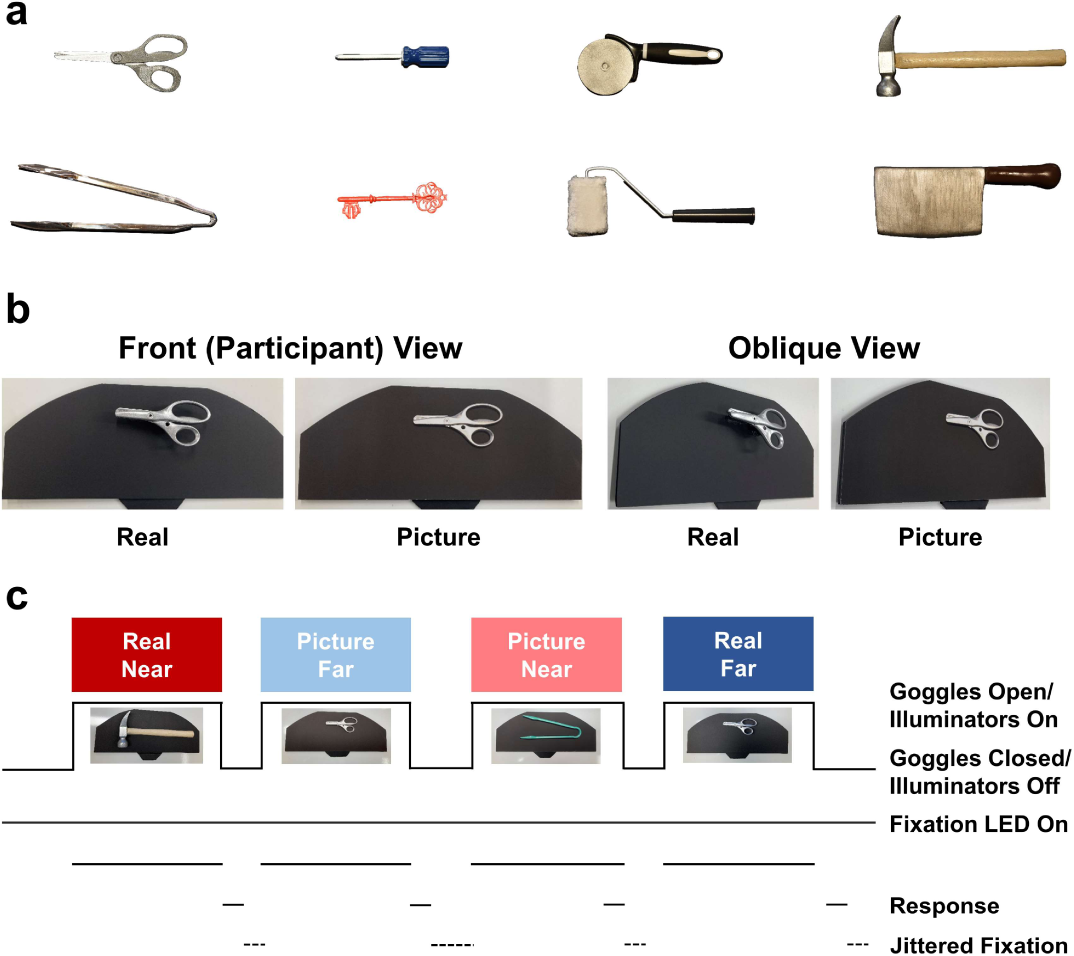
(a) Exemplars of the eight stimulus types used in the experiment. Each stimulus type was paired with a second stimulus type that required a similar hand action to use. Participants completed a 1-back task in which they evaluated whether the current tool uses the same hand action as the previous tool. (b) Front and oblique views of an example stimulus displayed as a real object and as a printed picture. (C) The trial sequence. In each of 64 trials per run, participants viewed stimuli for 2 seconds before making a button-press response with the left hand. Participants fixated on a red LED positioned above either the near or far stimulus location, in accordance with the placement of the stimulus for each trial.

The real object and picture stimuli were mounted on a large, angled, black wooden platform that was placed on the scanner bed over the participant’s waist (**Figure 1**). The stimuli were positioned into small slots cut into the midline of the platform. The ‘near’ slot was 13 cm from the front edge of the platform, while the ‘far’ slot was 51 cm from the front of the platform (38 cm between slots). For each participant, the platform was placed so that stimuli mounted in the near slot were within reach (typically ∼36 cm from the head), while those mounted in the far slot lay outside of reach.

Two red light-emitting-diodes (LEDs) served as fixation points. The fixation LEDs were located just above the stimulus, in equivalent positions relative to their respective stimuli in the near and far positions, thus ensuring that the retinal location of the stimuli was consistent across distance conditions. Fixation was individually calibrated for each participant. Both fixation points were calibrated to appear as a single fixation above the stimulus in both the near and far position. Critically, a growing body of behavioral research indicates that physical size is an especially important cue for recognizing real objects (Holler et al., 2019; Sensoy et al., 2021), so we kept the physical size of the stimuli the same across the near vs. far locations. Consequently, the same object subtended a larger retinal extent at the near position than at the far position. Bright white LEDs were used to illuminate the stimuli. Separate LEDs were used to illuminate the stimuli in the near and far locations, each with a similar angle of orientation towards the stimulus. The fixation and illuminator LEDs were held in place using separate strands of Loc-Line modular hose (Lockwood Products, Inc.). Two additional (green and red) low-illumination LED lights, which were affixed to the outside of the MRI bore and not visible to participants, served as guides for the experimenters to signal trial timing. Stimulus visibility was controlled on all trials using MRI-compatible liquid crystal glasses that were mounted to the front of the head coil. All trials were recorded for accuracy using an MR-compatible infrared bore camera (MRC Systems, GmbH) positioned behind the head coil.

### Task and Procedure

The participant’s task was to judge whether the currently-presented tool had a similar or different hand action to the tool shown on the previous trial. Before the experiment, participants completed a series of practice trials to ensure that they understood the task and the typical hand/arm action associations of the different objects. Participants entered their decision (similar or different) on each trial via button press on a response pad positioned under the left hand. Participants were instructed to maintain their gaze throughout all trials on the red fixation point that appeared just above the stimulus at the near or far location.

We tilted the head coil forwards by approximately 30° to allow participants to view the stimuli directly, without the use of mirrors. Stimulus viewing time was controlled using liquid crystal visual occlusion spectacles (PLATO, Translucent Technologies, Inc.) that switched between transparent (‘open’) and opaque (‘closed’) states. At the start of each trial, the fixation point and illuminator light at either the near or far location turned on, and the glasses opened to reveal the stimulus on the platform. The stimulus was visible for 2 seconds, after which time the glasses closed and the illuminator light turned off (fixation remained on for the duration of the trial). The glasses remained closed during the intertrial interval (ITI). During this time, participants made a response (within a 2-second time limit), and the experimenters changed out the stimuli on the platform. A team of four experimenters was required to quickly replace stimuli between trials and to efficiently sort stimuli in the order specified for the subsequent run. To decorrelate hemodynamic responses of successive trials, we used a jittered intertrial interval of either 2 or 4 seconds after the 2-second response period (**Figure 2c**). All runs had an equal number of 2- and 4-second ITIs. The distribution of ITIs was counterbalanced across conditions in each run. Event timing was controlled using custom MATLAB scripts (The Mathworks, Inc.). The bore light remained on throughout all scans. Aside from the sources of illumination inside the bore, and the small “experimenter” LEDs fixed to the outside of the magnet, the environment outside of the bore was otherwise dark. We used blackout material to cover the illuminated displays on the front of the scanner.

### Experimental Design and Statistical Analyses

The study had a 2 X 2 design with factors of Display Format (real objects vs. 2-D pictures) and Distance (near vs. far). Each combination of the 16 tool exemplars (8 tool types X 2 exemplars of each) X 2 display formats X 2 distance conditions (64 trials) was presented once per run. The order of trials in each condition was pseudorandomly determined in each run, with the constraint that the same tool was not presented on consecutive trials. Each run began and ended with 12 and 8 seconds respectively of resting baseline fixation. Each run lasted 468 seconds, and the entire experiment lasted ∼2 hours in total, including set-up time. Depending on time constraints, most participants completed 8 experimental runs. In addition to the main experimental runs, an anatomical scan was also collected for each participant.

### Data Acquisition

All data were collected at Renown Regional Medical Center (Reno, Nevada) using a 3-tesla Phillips Ingenia scanner with a 32-channel digital SENSE head coil. Functional data were collected using a T2*-weighted single-shot gradient-echo echo-planar imaging (EPI) sequence [time to repetition (TR) = 2,000 ms; time to echo (TE) = 30 ms; slice thickness = 3 mm; field of view = 240 x 240 mm; matrix size = 80 x 80 pixels; flip angle = 76°]. Each functional volume was comprised of 40 oblique slices that were acquired with a ∼30° caudal tilt with respect to the plane of the anterior-posterior commissures, providing near whole-brain coverage. The anatomical image was collected using a 3-D T1-weighted pulse sequence (TR = 3000 ms; TE = 4.6 ms; matrix size 256 x 256 mm; flip angle = 8°; 1mm isotropic voxels).

### Data Analysis Software

Analyses were performed using BrainVoyager QX 3.6 (Brain Innovation, B.V.), NeuroElf (https://neuroelf.net/), The Representational Similarity Toolbox (Nili et al., 2014), and custom code written in MATLAB (The MathWorks, Inc.).

### FMRI Data Preprocessing

Functional data preprocessing included slice scan-time correction (cubic spline interpolation for slices acquired in ascending interleaved order), 3-D motion correction (trilinear/sinc interpolation), and high-pass GLM-Fourier temporal filtering (cutoff of 2 sine/cosine cycles per run). Anatomical data were normalized to MNI-ICBM 152 space following an iso-voxel spatial transformation (trilinear interpolation) and intensity inhomogeneity correction (3 cycles). Each functional run was separately co-registered to the anatomical scan. To improve signal quality, functional data were denoised using GLMdenoise (Kay et al., 2013). Runs were excluded in cases where 3-D motion correction parameters indicated excessive head movement (12 runs), and in cases where screening of the offline video recordings revealed errors with stimulus presentation or trial timing (9 runs). For the univariate analyses, functional data were spatially smoothed using a 6-mm full width at half maximum Gaussian filter. Because pattern-based analyses benefit from finely differentiated spatial information, spatial smoothing was not used in the preprocessing pipeline for the multivariate representational similarity analysis (RSA).

### General Linear Models

Beta weights were calculated for each condition using a random-effects general linear model (GLM) that included a predictor for each condition convolved with BrainVoyager’s ‘two-gamma’ default hemodynamic response function and aligned to trial onset. For the univariate analyses, the GLM included 4 predictors (Real Near, Real Far, Picture Near, Picture Far). For the multivariate analyses, the GLM had 32 predictors, one for each of the eight stimulus identities presented as each Display Format (Real, Picture) and in each Location (Near, Far). Event-related averages were calculated using a baseline extending from 2 seconds before trial onset and 12 seconds after trial onset.

### Univariate data analysis

We first examined whether there were brain areas that responded more strongly to real objects than pictures (and to pictures more strongly than real objects) by searching for a main effect of Display Format using the contrasts ([+Real Near +Real Far] > [+Picture Near +Picture Far]), and ([+Picture Near +Picture Far] > [+Real Near +Real Far]), respectively. Next, we examined whether there was a main effect of Distance by searching for areas in which activation was significantly greater for near than far stimuli (and vice versa) using the contrasts ([+Real Near +Picture Near] > [+Real Far +Picture Far]) and ([+Real Far +Picture Far] > [+Real Near +Picture Near]), respectively. Finally, we tested the interaction between Distance and Display Format using the contrasts (a) ([+Real Near +Picture Far] > [+Real Far +Picture Near]) to identify brain areas for which a near location amplified brain responses more strongly for real objects than pictures, and (b) ([+Real Far +Picture Near] > [+Real Near +Picture Far]) to identify brain areas for which a near location amplifies brain responses more strongly for pictures than real objects. For each contrast, the resultant group activation maps were set to a minimum statistical threshold of *p* < 0.001.

### Definition of Regions of Interest

Multivariate analyses were performed in regions of interest that included early visual cortex (EVC), areas within the ventral and dorsal streams thought to be involved in visual recognition and visually-guided action, and somatomotor areas. ROIs were defined based on the univariate contrast of response versus baseline (false discovery rate < .05), which elicited widespread activation throughout early visual cortex, the ventral and dorsal visual streams, as well as motor and somatosensory cortex. EVC was selected for its role in early visual processing. We selected dorsal regions SPOC, pIPS, aIPS, S1, M1, and PMd because of their involvement in the fronto-parietal reaching-and-grasping network (Gallivan et al., 2013; Gallivan and Culham, 2015; Rens et al., 2023). LOC was included for its role in object shape processing (Malach et al., 1995; Grill-Spector et al., 2001; Kourtzi and Kanwisher, 2001), and its differential activity in response to repetition of real objects vs. pictures (Snow et al., 2011). pMTG was included for its role in visually guided action and mental imagery of tool use (Johnson-Frey et al., 2005; Fabbri et al., 2016). VMT was included because of its putative role in haptic shape processing (Snow et al., 2014).

We focused our investigation on data derived from left hemisphere ROIs because of their relationship to the dominant hand of our (right-handed) participants. Analyses in corresponding right-hemisphere ROIs, where they were available, revealed similar patterns of results as in the left hemisphere (see **Supplementary Figures S1-5 and Supplementary Tables S1-S5**). ROIs were defined separately for each participant, and independently from the subsequent RSA analysis to avoid any selection bias (Kriegeskorte et al., 2008). Regions were defined based on anatomical and activation criteria rather than stereotaxic coordinates. ROI centers were localized by starting with the point of maximum activation nearest to expected anatomical landmarks (see **Table 1** for mean left-hemisphere ROI centers and **Supplementary Table S1** for mean right-hemisphere ROI centers). In some cases, the ROI center was shifted slightly to avoid overlap with other nearby ROIs, sulcal boundaries, or cortical edges. ROIs consisted of the active voxels within a cube with side length equal to 10 units of BrainVoyager’s internal coordinate system, roughly equivalent to 10X10X10 MNI coordinates, or 64 functional voxels.

**Table 1.** MNI coordinates for left-hemisphere regions of interest used in multivariate analyses.

| ROI | Mean x-coordinate | Mean y-coordinate | Mean z-coordinate |
| --- | --- | --- | --- |
| EVC | -7 | -89 | 2 |
| LOC | -42 | -74 | -11 |
| pMTG | -46 | -66 | -8 |
| VMT | -23 | -61 | -14 |
| SPOC | -10 | -77 | 38 |
| plPS | -27 | -67 | 40 |
| alPS | -39 | -39 | 51 |
| S1 | -40 | -29 | 55 |
| M1 | -35 | -21 | 52 |
| PMd | -28 | -3 | 54 |

The regions were localized using the following landmarks: early visual cortex (EVC) was localized at the site of peak activation near the calcarine sulcus (Snow et al., 2014); lateral occipital complex (LOC) was localized at the site of peak activation in the vicinity of the junction between the inferior temporal sulcus and the lateral occipital sulcus (Snow et al., 2015); posterior medial temporal gyrus (pMTG) was localized towards the posterior end of the medial temporal gyrus, with MNI coordinates from a previous study used as a reference point (Fabbri et al., 2016); ventromedial temporal cortex (VMT) was localized immediately dorsal to the cerebellum, with particular attention to matching bilateral position between hemispheres (Snow et al., 2014); superior parieto-occipital cortex (SPOC) was localized at the site of peak activation near the superior end of the parieto-occipital sulcus (Gallivan et al., 2009); posterior intraparietal sulcus (pIPS) was localized at the site of peak activation near the posterior end of the intraparietal sulcus (Roth and Zohary, 2015; Fabbri et al., 2016); anterior intraparietal sulcus (aIPS) was localized at the site of peak activation near the anterior end of the intraparietal sulcus, at its junction with the postcentral sulcus (Cavina-Pratesi et al., 2010; Fabbri et al., 2016); primary somatosensory cortex (S1) was localized at the site of peak activation on the dorsal postcentral gyrus (Case et al., 2016); primary motor cortex (M1) was localized on the hand knob of the precentral gyrus (Yousry et al., 1997); and dorsal premotor cortex (PMd) was localized at the site of peak activation around the junction of the superior frontal sulcus and the precentral sulcus (Wang et al., 2008).

### Representational Similarity Analysis (RSA)

The Representational Similarity Toolbox (MATLAB version) was used for RSA. Beta estimates for each of the 32 conditions (8 tool types X 2 display formats X 2 distance conditions) were extracted at each voxel using non-smoothed, denoised, single-subject GLMs calculated across runs. Within each ROI, for each pairwise comparison of the 32 conditions, one minus the Pearson’s correlation coefficient was calculated across beta estimates for all voxels, yielding a representational dissimilarity matrix (RDM) for each subject. The RDMs for all subjects were averaged to create a group-level RDM, which was used for multidimensional scaling (MDS) analyses. Individual-subject RDMs were used to estimate performance of representational models and to calculate noise ceiling estimates.

Representational models based on Distance, Display Format, Grip Type, Stimulus Identity, and Size were compared with the data RDMs. Each of the model RDMs represents a hypothesis about what stimulus features are most important in determining the representational geometry within an ROI. The Distance model assumes that stimuli placed within reach are represented similarly to each other and differently from stimuli placed outside of reach, the latter of which are also represented similarly to each other. Likewise, the Display Format model assumes that the prevailing factor for stimulus representation is whether stimuli are presented as real objects or as pictures. The Grip Type model assumes that the prevailing representational factor is whether the exemplar object is typically grasped with a precision grip focused on the fingertips or grasped with a power grip using the whole hand. The Identity model assumes that only stimuli of the same tool type are represented similarly. The Size model categorically groups the four largest stimulus types together (cleaver, hammer, tongs, paint roller) and the four smallest stimulus types together (key, scissors, pizza cutter, screwdriver). We also tested a model based on the hand action pairs, but these data are not considered further because the shared representational geometry of the Hand Action and Grip Type models led to redundant patterns of results.

The performance of each model was assessed using the Spearman’s Rank Correlation Coefficient (Kriegeskorte et al., 2008) comparing each model to each subject’s RDM, subsequently averaged across participants. T-tests determined whether the correlation was significantly greater than zero. Statistical significance was Bonferroni-corrected from α=.05 for the number of models tested within each region. Results that were significant prior to but not following Bonferroni correction are noted as marginally significant. Averages of data-model correlations across ventral ROIs and across dorsal ROIs were also calculated. Within each ROI, noise ceiling estimates were calculated to assess the between-subjects variability among the RDMs. The lower bound of the noise ceiling was derived using a leave-one-out cross-validation approach, while the upper bound of the noise ceiling represents the performance of a hypothetical model constructed to maximize the average correlation with all of the individual subjects’ RDMs (Nili et al., 2014).

While we focused our analyses on the theoretical models described above, they are not exhaustive of all possible stimulus characteristics that could describe the representations. For example, we did not include models of lower-level stimulus characteristics such as luminance and contrast (Bohon et al., 2016), and we did not explicitly test models that related to other higher-level physical object characteristics such as surface texture or weight, since some of our stimuli were artificial (for compatibility with high-field MR environments), and as such they were not perfectly representative of typical real-world physical object characteristics. Furthermore, while we have focused our analyses on orthogonal models that test the contribution of unique stimulus characteristics to the representations, the models we tested may be combined in numerous ways to examine more complex interaction effects.

Initial representational similarity analyses revealed that the Distance model had a powerful effect on fMRI response patterns across all ROIs (see *Egocentric distance is a principal organizing factor of object representations*). In fact, the Distance model swamped the effects of all of the other models, making their performance difficult to evaluate. As such, to gauge the relative performance of the remaining models more effectively, we separated the data between trials in the Near vs. Far conditions and conducted RSA separately on the two datasets (see *Egocentric distance modulates object representations in dorsal more than ventral cortex)*. We also used MDS to visualize the representations in each ROI (see *Multidimensional Scaling*). For the MDS analysis of stimulus size, the data were further separated according to the display format of the stimulus (see *Size representations depend on a combination of distance and display format in dorsal but not ventral cortex*).

### Multidimensional Scaling

The group-averaged representational dissimilarity matrices were used as the input for MDS (Torgerson, 1958; Kruskal, 1964; Borg and Groenen, 2005), in which data points representing the response in an ROI to each stimulus are placed in a multidimensional space where increasing distances in the space represent increasing dissimilarity between responses. All MDS analyses were restricted to two-dimensional spaces for ease of visualization. Metric stress was used as the goodness-of-fit criterion. To test whether points in the MDS space clustered according to identifiable stimulus categories, we used t-tests to compare the pairwise differences for stimuli within the same category against the pairwise distances for stimuli of different categories. Statistical significance was Bonferroni-corrected from α=.05 according to the number of category types tested within each region. As above, results that were significant prior to but not following Bonferroni correction are noted as marginally significant.

## Results

### Display format and egocentric distance modulate fMRI BOLD amplitudes

To explore how egocentric distance and display format influence fMRI BOLD response amplitudes, we computed a group voxelwise random-effects GLM and contrasted responses to real objects vs. pictures (a main effect of Display Format), and to near vs. far stimuli (a main effect of Distance). We also tested for a Display Format X Distance interaction to determine whether distance influenced fMRI amplitude responses differently between the two display formats. Because we used an event-related design in which trials of each Display Format and Distance condition were randomly interleaved throughout the experiment, participants did not know from trial to trial what the upcoming stimulus would be or where it would be located, thus minimizing the likelihood that differences could be attributed to anticipatory effects (Clark, 2012). Brain responses for the main effect and interaction contrasts are displayed in **Figure 3** on representative slices of a group-averaged anatomical image. We applied a threshold of *p* < 0.001 (uncorrected) for all contrasts to ensure that areas of activation reflect meaningful differences, and to demonstrate than an absence of (otherwise meaningful) activation cannot be attributable to arbitrary thresholding decisions (Snow et al., 2015). MNI coordinates of activated regions in each contrast, together with the associated cluster sizes and *t*-statistics, are presented in **Table 2**.

**Figure 3:**
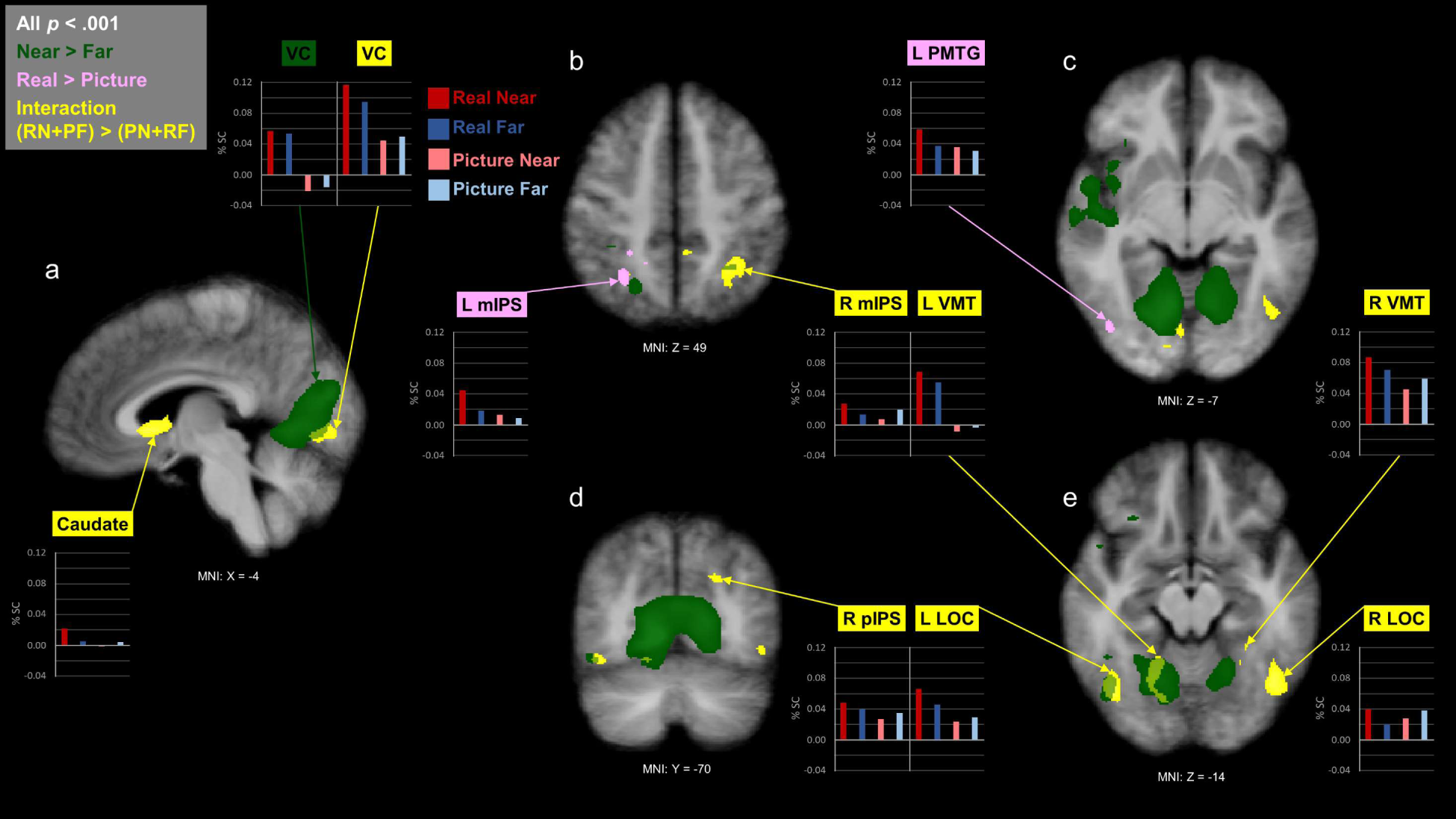
Univariate contrasts displayed on sagittal (a), axial (b,c,e) and coronal (d) slices of the group-averaged brain. Pink shading shows voxels where stimuli presented as Real Objects elicited stronger activation than stimuli presented as Pictures. Green shading shows voxels where stimuli in the Near condition elicited stronger activation than stimuli in the Far condition. Yellow shading shows voxels where the difference in activation between the Near and Far conditions is greater for Real Objects than for Pictures. All contrasts, threshold of p<.001. For illustrative purposes, in the clusters identified by the contrasts described above, inset bar charts show peak mean % SC from resting baseline for stimuli in the Real Near, Real Far, Picture Near and Picture Far conditions. VC (visual cortex); L and R mIPS (left and right middle intraparietal sulcus); R pIPS (right posterior intraparietal sulcus); L pMTG (left posterior medial temporal gyrus); L and R VMT (left and right ventromedial temporal cortex); L and R LOC (left and right lateral occipital complex); caudate (head of caudate nucleus).

**Table 2:** MNI coordinates, size in functional voxels, and mean *t*-statistic for clusters identified in the univariate contrasts.

| Contrast | ROI | x-coordinate | y-coordinate | z-coordinate | Size in functional voxels | Mean *t*-statistic |
| --- | --- | --- | --- | --- | --- | --- |
| **Main effect of display format, Real > Picture**<br>[(+Real Near +Real Far) > (Picture Near +Picture Far)]<br>Clusters>100 voxels | Left IPS | -27 | -57 | 49 | 215 | 4.231 |
|  | Left LOC/pMTG | -42 | -75 | -8 | 137 | 4.285 |
|  | Left IPS | -24 | -43 | 46 | 123 | 4.095 |
| **Main effect of display format, Picture > Real**<br>[(+Picture Near +Picture Far) > (+Real Near +Real Far)]<br>Cluster>100 voxels | Right occipital | 22 | -94 | -13 | 280 | -4.311 |
| **Main effect of distance, Near > Far**<br>[(+Real Near +Picture Near) > (+Real Far +Picture Far)]<br>Clusters > 400 voxels | Primary visual | -12 | -73 | -2 | 42871 | 5.217 |
|  | Left insula | -42 | -31 | 12 | 9923 | 4.533 |
|  | Left anterior temporal | -45 | -1 | -41 | 2171 | 4.278 |
|  | Left frontal | -52 | 32 | 25 | 547 | 4.425 |
|  | Left IPS | -21 | -63 | 49 | 436 | 4.147 |
|  | Left LOC | -42 | -61 | -11 | 434 | 4.241 |
| **Main effect of distance, Far > Near**<br>[(+Real Far +Picture Far) > (+Real Near +Picture Near)]<br>Clusters > 400 voxels | Left occipital pole | -22 | -100 | -8 | 9079 | -4.81 |
|  | Right occipital pole | 33 | -97 | -11 | 8633 | -4.93 |
|  | Right parietal | 39 | -78 | 37 | 5465 | -4.418 |
| **Interaction contrast**<br>[(+Real Near +Picture Far) > (+Picture Near +Real Far)]<br>Clusters > 300 voxels | Caudate | -3 | 15 | -2 | 1709 | 4.744 |
|  | Right LOC | 45 | -62 | -14 | 1361 | 4.757 |
|  | Left LOC | -42 | -67 | -14 | 799 | 4.531 |
|  | Right IPS | 32 | -52 | 47 | 687 | 4.457 |
|  | Left VMT | -18 | -55 | -12 | 654 | 4.42 |
|  | Visual cortex | -3 | -75 | 5 | 633 | 4.378 |
|  | Left occipital | -24 | -84 | 16 | 302 | 4.681 |
| **Reverse Interaction contrast**<br>[(+Picture Near +Real Far) > (+Real Near +Picture Far)]<br>Clusters > 100 voxels | Dorsal cerebellum | -5 | -46 | -2 | 156 | -4.389 |

If a brain area responds more strongly to real objects than to pictures irrespective of stimulus position, it should show stronger responses to real objects presented at either distance (Real Near and Real Far) versus pictures presented at either distance (Picture Near and Picture Far). Accordingly, using the contrast [(+Real Near +Real Far) > (+Picture Near +Picture Far)], we found that real objects elicited stronger responses than pictures in the vicinity of the lateral occipital complex (LOC) and posterior middle temporal gyrus (pMTG) in the ventral visual pathway, and in the intraparietal sulcus (IPS) of the dorsal visual pathway (**Figure 3**, pink). Both LOC and IPS have previously been implicated in shape-related processing (Konen and Kastner, 2008; Orban, 2011; Freud et al., 2017). Notably, the reverse contrast, which identified brain areas that showed stronger responses to pictures at either distance than to real objects at either distance [(+Picture Near +Picture Far) > (+Real Near +Real Far)], revealed only one sizable cluster of significant activation in right occipital cortex. These results indicate that real objects drive brain responses more strongly than pictures do in higher-level temporal and parietal cortical areas commonly associated with shape processing.

Next, we tested whether the egocentric distance of our stimuli modulated fMRI BOLD responses. A brain area that responds more strongly when a stimulus is closer to the observer than when it is farther away should show stronger responses to the near (Real Near and Picture Near) versus far (Real Far and Picture Far) stimuli in our study, irrespective of format. The contrast [(+Real Near +Picture Near) > (+Real Far + Picture Far)] revealed comparatively large swaths of activation across the brain (**Figure 3**, green), with the largest clusters in the vicinity of early visual cortex, left insula, and left anterior temporal cortex. Sizable clusters of activation from this contrast were also observed in left frontal cortex, left IPS, and left LOC. The reverse contrast, [(+Real Far +Picture Far) > (+Real Near +Picture Near)], which indicates stronger activation for far than for near stimuli, revealed comparatively less activation, with small clusters limited to the left and right occipital poles, and one in right posterior parietal cortex.

We kept the physical size of our stimuli constant across the near and far egocentric distances because the physical size of everyday, familiar real objects, such as those used in the current study, provides a salient cue to identity. For example, when the size of real objects is manipulated so that they deviate from their typical real-world size, recognition is disrupted, whereas analogous size manipulations do not influence recognition of pictures of objects (Holler et al., 2019; Sensoy et al., 2021). Therefore, our design ensured that recognition processes would be comparable across formats. Of course, if brain regions show stronger responses to stimuli in the Near than the Far condition, as did many areas in the Distance main effect contrast described above, this could reflect either a sensitivity to egocentric proximity, or a sensitivity to retinal extent via the activity of cortical visual field maps (Bandettini et al., 1992; Kwong et al., 1992; Ogawa et al., 1992)–since when the same object is positioned closer to (vs. farther from) the observer, it occupies a larger part of the visual field and thus activates adjacent parts of the cortex. Such retinotopic maps have been identified not only in early occipital cortical areas, but also along ventral temporal cortex (including area LO) and dorsally in the intraparietal sulcus and superior parietal lobule (SPL) (see Wandell & Winawer, 2011 for review). It is perhaps because of the widespread cortical sensitivity to retinal extent that most fMRI studies of object vision use scaled stimuli that subtend the same retinal angle at the same viewing distance, even though their typical sizes differ by orders of magnitude in the real world (Snow and Culham, 2021). Nevertheless, if real and picture stimuli are differentially influenced by distance, this would suggest that the distance effects on BOLD amplitudes cannot be fully explained by retinal extent, since the size of the stimuli in each format are identical at the near and far locations.

To determine whether distance influenced fMRI amplitude responses differently between the two display formats, we tested for an interaction between Format and Distance using the contrast [(+Real Near +Picture Far) > (+Picture Near +Real Far)]. Because previous electrophysiological studies have reported that brain responses to real objects are amplified in dorsal cortex compared to pictures (Marini et al., 2019), especially when the stimuli are within reach and accessible for interaction (Fairchild et al., 2021), we predicted that nearness would increase activation more for real objects than for pictures, particularly in dorsal cortex (Freud et al., 2020; Snow and Culham, 2021). The interaction contrast produced significant activation, which was typically bilateral, in dorsal areas including middle and posterior IPS, as well as ventrally in areas including LOC and ventromedial temporal cortex (**Figure 3**, yellow). Notably, many of these areas overlapped with regions activated in the Distance main effect contrast. A test of the opposite interaction—to identify brain areas in which nearness increased activation more strongly for pictures than for real objects—using the contrast [(+Picture Near +Real Far) > (+Real Near + Picture Far)] revealed little, if any, significant activation. To investigate this pattern further, we computed the mean peak percent signal change (% SC) above resting baseline for our four experimental conditions within brain regions activated in the interaction contrast, as well as for other areas of activation revealed by the main effect contrasts, as displayed in **Figure 3**. We found that in all areas, nearby real objects elicited qualitatively higher responses than any of the other three conditions, indicating that in these regions, fMRI BOLD responses tended to be stronger for nearby real objects than for distant real objects and for pictures presented in either distance condition. Taken together, these data indicate that BOLD amplitudes to real objects were boosted when the stimulus was positioned at a near vs. far egocentric distance, whereas this was not the case for pictures, for which nearness had little effect on responses.

### Multivariate analyses

Previous behavioral work has shown that observers categorize real object stimuli according to a wider variety of stimulus attributes than pictures, including physical attributes such as size (Holler et al., 2020). Although one previous fMRI study compared BOLD responses during perception of real objects versus pictures (Snow et al., 2011), that study used a slow event-related design that had too few trials for multivariate analyses. Therefore, we used multivariate pattern analysis to examine the relative influence of display format, egocentric distance, and stimulus size on object representations. Whereas our univariate analyses averaged across trials in which different objects were presented in the same condition (e.g., near real stimuli), our multivariate analyses take advantage of the variability among the objects to reveal how the brain represents different object features at the level of individual object types (de-Wit et al., 2016). Moreover, while univariate effects are assessed separately at each voxel, multivariate analyses compare patterns of activity across voxels—thereby improving sensitivity (because meaningful patterns may emerge across voxels) and maximizing statistical power (because multiple comparisons corrections are not applied for separate tests across thousands of voxels) (Cremers et al., 2017).

For the multivariate analyses described below, we used representational similarity analysis (RSA) (Kriegeskorte et al., 2008) to compare response patterns in key cortical regions to theoretical models of which stimulus features might be represented in these regions. We also used multidimensional scaling (MDS) (Kruskal, 1964) to visualize the similarities and differences among patterns of cortical representation for the stimuli used in the experiment. MDS organizes fMRI responses to each stimulus in a multidimensional space, such that stimuli that elicit similar response patterns across voxels within an ROI are placed nearer to one another than to stimuli that elicit dissimilar response patterns. Apparent groupings between items in the MDS plots arise due to the underlying similarity in the cortical representations of those stimuli. Because the positioning of items within the representational space is unconstrained by *a priori* theoretical models, MDS allows a unique, data-driven visualization of the similarities and dissimilarities between the representations that is distinct from that conveyed by the data-model correlations. We also tested *a posteriori* whether stimulus groupings in the multidimensional space for each ROI corresponded to particular characteristics, such as format, distance, or size (see **Tables 3-6**).

Analysis of the entire dataset using RSA and MDS showed a powerful effect of egocentric distance on cortical response patterns. Specifically, stimuli presented at the near position elicited highly distinct neural representations compared to those at the far position. In fact, egocentric distance swamped the effects of all other models in the representational similarity analyses, and the effects of distance were also clearly visible in model-free MDS arrangements. Therefore, to explore the extent to which other stimulus features influenced representations over and above egocentric distance, we split the data between the two egocentric distances and conducted RSA and MDS analyses of object representations separately at the near and far stimulus locations. Finally, because accumulating evidence from behavior and neuroimaging studies suggest that stimulus size is processed more readily in the case of real objects than of pictures (Holler et al., 2019; Sensoy et al., 2021), and that stimulus distance modulates responses to real objects (Gallivan et al., 2009, 2011; Cavina-Pratesi et al., 2010) but less so for pictures (Gomez et al., 2018; Snow et al., 2023), we further split the data according to display format to examine whether stimulus size influenced representational patterns differently according to the combination of display format and egocentric location. We focus here on ROIs defined in the left hemisphere, consistent with the hemisphere of motoric dominance of our right-handed participants; however, the pattern of results in the right hemisphere (**Supplementary Figures S1-5**) was strikingly similar to that in the left.

### Egocentric distance is a principal organizing factor of object representations

We began by using RSA to examine whether stimulus characteristics such as egocentric distance, identity, and size were represented in the fMRI object response patterns. We were also interested in whether cortical representations would reflect the format in which the stimulus was displayed. We tested fMRI response patterns derived from key ventral and dorsal ROIs against different theoretical models, each of which instantiated a hypothesis about the stimulus characteristics that govern the representations. We computed a data representational dissimilarity matrix (RDM) for each observer that measured dissimilarity between fMRI responses to each stimulus in each condition. **Figure 4a** shows an example RDM derived from fMRI responses in LOC; for illustrative purposes, the RDM represents data that has been averaged across subjects. **Figure 4b** shows the theoretical models against which the data RDMs for each subject were tested. We tested five orthogonal models: Distance, Display Format, Grip Type, Identity, and Size (see *Materials and Methods*). Each model assumes that stimuli that share a particular characteristic are represented similarly to each other, and dissimilarly to stimuli that differ along the relevant characteristic; the influence of all other stimulus characteristics on the representations are held constant in the model. For example, the Distance model assumes that all near stimuli are represented similarly, all far stimuli are represented similarly, and that near and far stimuli are represented differently from each other.

**Figure 4.**
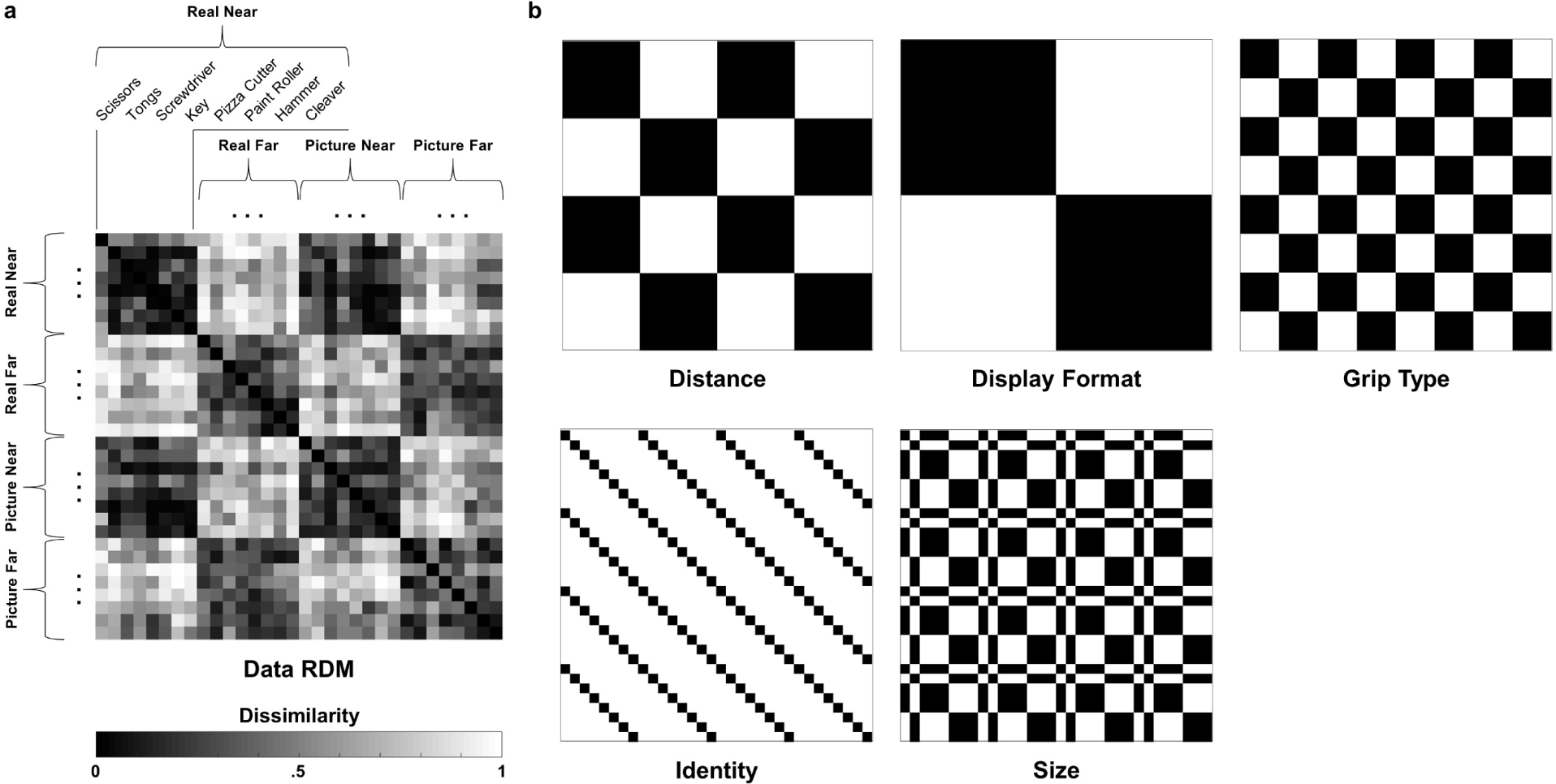
(a) An example data RDM derived from fMRI responses in LOC. For illustrative purposes the RDM shows data that has been averaged across subjects; however, the RSA analyses were conducted iteratively using the single subject RDMs. Each cell in the matrix of the data RDM represents the similarity of fMRI responses across voxels in the ROI to a pair of stimuli presented during the study. The 32 object exemplars used in our experiment are ordered along the horizontal and vertical axes of the matrix according to display format and distance, and items appear in the same order along both axes; thus, the upper right and lower left triangles of the matrix are identical. (b) Theoretical model RDMs against which fMRI responses (data RDMs) in each ROI were tested. Each model RDM represents a hypothesis about which stimulus features are represented in fMRI response patterns in each ROI.

The mean Spearman’s Rank Correlation between the data RDMs and each theoretical model is displayed in **Figure 5**, separately for each ROI. MDS plots for each ROI are shown in **Figure 6**. As a measure of the maximal model performance attainable, the data-model correlations shown in **Figure 5** display the upper and lower bound estimates of the noise ceiling, separately for each ROI. The noise ceiling estimates reflect the fact that a single RDM model is unlikely to correlate perfectly with the data RDMs, because a given brain region is unlikely to be selective for just one stimulus feature, and because there is inherent noise in fMRI data due to signal quality and variability between participants. For any given model, the distance between the noise ceiling and unity is proportional to the noise in the data (i.e., higher noise in the data corresponds to a lower noise ceiling), while the distance between model performance and the noise ceiling is proportional to the signal in the data left unexplained by the model (i.e., better-performing models approach the noise ceiling more closely). We found that noise ceilings tended to be higher in ventral (vs. dorsal) ROIs, indicating that neural representations in ventral cortex were more consistent across observers, dovetailing with findings from previous studies (Bracci et al., 2017; Bracci & Op de Beeck, 2016; Fabbri et al., 2016). Because of the variability in noise ceilings between ROIs, we focused our analyses on model performance within each ROI, rather than between ROIs (Fabbri et al., 2016).

**Figure 5:**
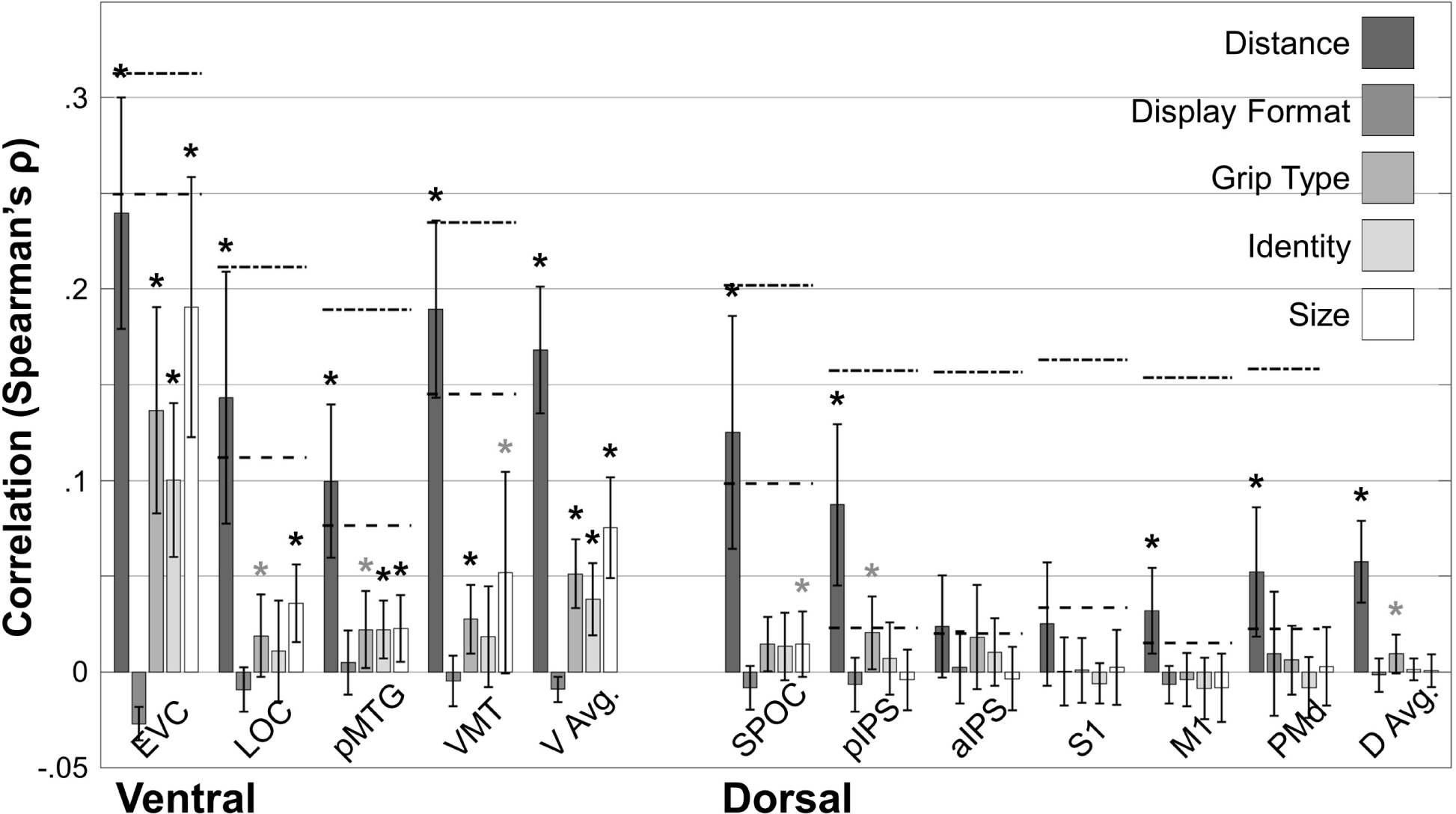
Spearman’s rank correlation coefficient, averaged across subjects, between each model RDM and the data RDMs in each ROI. A stronger correlation indicates that a model better describes the features represented in a given ROI. Black asterisks indicate correlations significantly greater than zero, while grey asterisks indicate marginal significance. The upper and lower bounds of the noise ceilings are shown with dashed lines. The noise ceiling is an estimate of how well a model could do if it were capturing all of the stimulus features an ROI represents, given the noise present in the data caused by factors unrelated to stimulus features.

**Figure 6:**
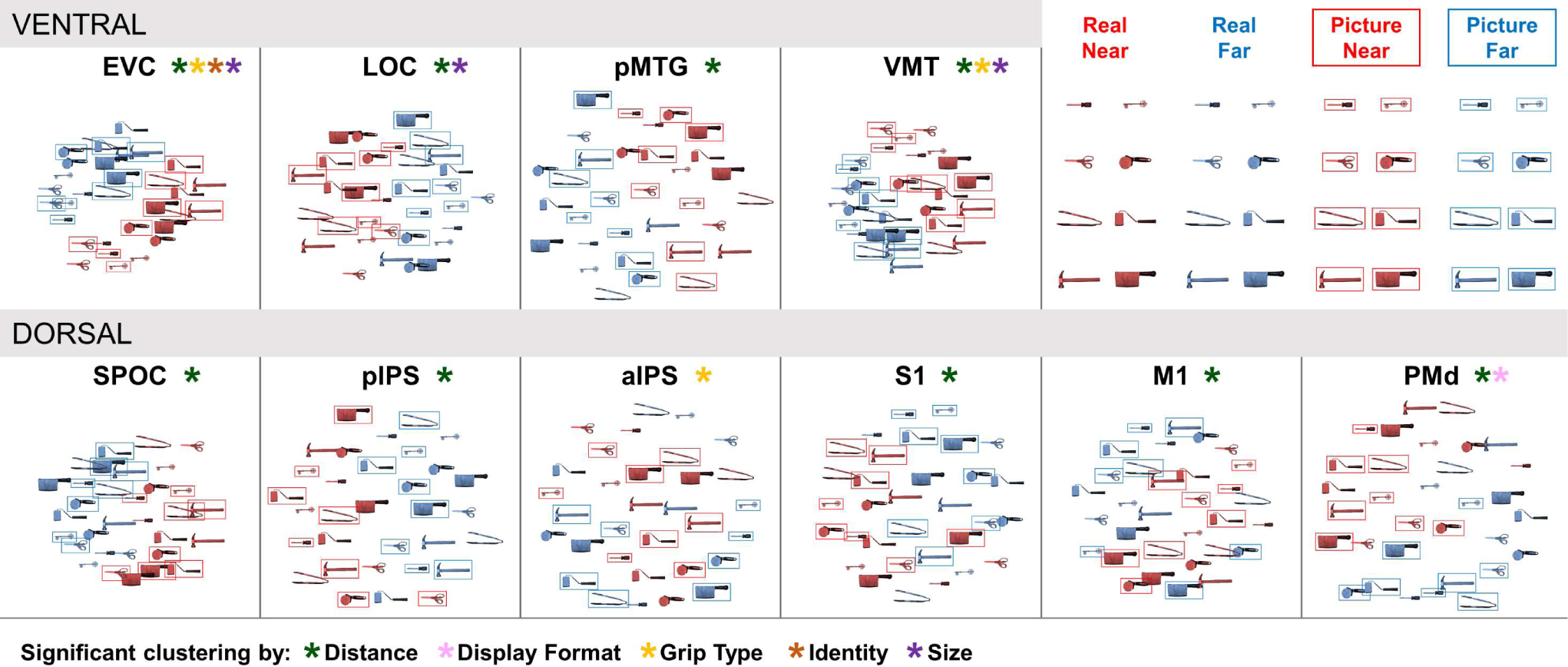
MDS plots for each ROI, ordered left-to-right along the posterior-anterior axis. Greater distance between a pair of stimuli in each MDS space corresponds to greater dissimilarity in activation patterns elicited by those two stimuli. For visualization purposes, stimuli in the Near condition are shown in red, while stimuli in the Far condition are shown in blue. Stimuli presented as pictures are depicted in rectangular frames, while stimuli presented as real objects are shown without such frames.

Consistent with the widespread sensitivity to egocentric distance that we observed in fMRI amplitudes in the univariate analyses, both the data-model correlations and the MDS plots evinced a clear organization of representations according to stimulus distance. In the data-model correlations, the Distance model was the only one to consistently approach the maximum model performance possible given the noise in the data, surpassing the lower-bound noise ceiling estimate in most ROIs. T-tests against zero confirmed that the Distance model performed significantly better than chance in EVC (p=4.95E-8), LOC (p=.000105), pMTG (p=2.52E-5), VMT (p=2.92E-8), SPOC (p=.000193), pIPS (p=.000177), M1 (.00387), and PMd (.00219). The Distance model did not reach significance in aIPS (p=.0389) and S1 (p=.0602), but it was nevertheless one of the strongest of the models in those areas. Distance also emerged as the principal organizing factor in the model-free MDS visualizations. As is apparent in **Figure 6**, near stimuli (illustrated in red) clustered separately from far stimuli (illustrated in blue). In ventral ROIs, the clustering by egocentric distance was so distinct as to be linearly separable. Clustering by distance in the MDS plots was significant in all ROIs except aIPS (**Table 3**). In summary, in both the data-model correlations and the MDS arrangements, distance predominated as the major organizing principle in most ROIs, and this was especially so in ventral cortex. The strong performance of the Distance model relative to the noise ceiling in each ROI suggests that the model accounted for a sizeable proportion of the observable signal. Finally, the clear organization of representations according to distance in the MDS plots indicates that the strong performance of the Distance model in the data-model correlations is unlikely to hide other, potentially better-performing theoretical models.

**Table 3:**
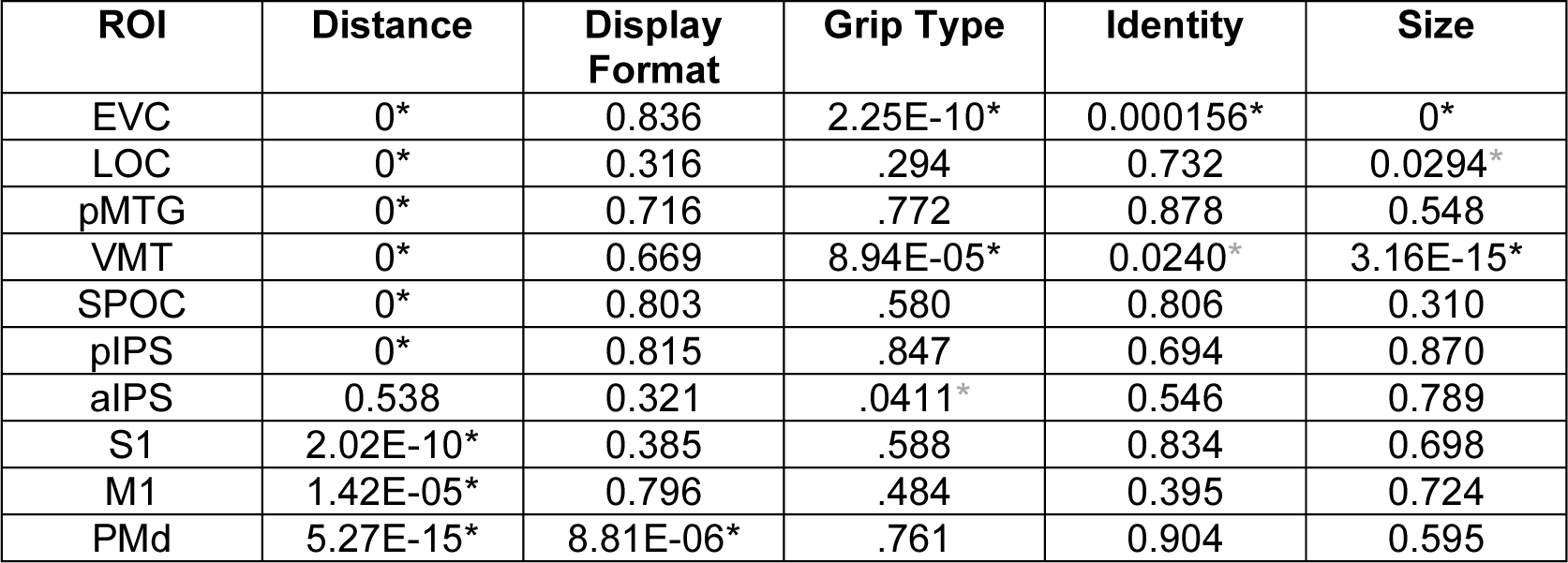
*P*-values for clusters in the MDS plots of Figure 6. *P*-values below 1E-15 are reported as 0. Here and in all other tables, significant results are marked with a black asterisk, while marginally significant results are marked with a gray asterisk.

Although multiple brain regions responded more strongly to real objects than to pictures in the univariate analyses, the Display Format model did not correlate significantly with the data RDMs in any ROI. Similarly in the MDS plots, representations did not cluster by Display Format in any of the ROIs, except for PMd (**Table 3**), for which representations of real objects clustered separately from those of pictures in a direction approximately orthogonal to the clustering of near vs. far stimuli. Taken together, these results suggest that while real objects (compared to pictures) boosted the overall strength of the BOLD response in some brain regions, a model accounting only for display format rarely explained the response patterns.

Next, we examined how identity, grip type, and size influenced response patterns. In the data-model correlations, the Identity model performed significantly above chance in EVC (p=2.49E-5) and pMTG (p=.00327). Clustering by identity was also observed in the MDS plots in EVC and marginally in VMT (**Table 3**). In LOC, identity was not significantly represented in either the data-model correlations or MDS plots, as might be expected from previous work suggesting that LOC codes object shape rather than object identity per se (Margalit et al., 2016; Freud et al., 2017). The Grip Type model was significant in EVC (p=1.95E-5) and VMT (p=.00234), and marginally significant in LOC (p=.0400) and pMTG (p=.0167), but it failed to reach significance in aIPS (p=.089), despite the putative role of this region in pre-shaping the hand for grasping (Gallese et al., 1994; Tunik et al., 2005). However, representations in the MDS plots did cluster marginally according to Grip Type in aIPS, as well as significantly in EVC and VMT (**Table 3**). Finally, yet perhaps most surprisingly, we found that stimulus size had a powerful influence on object representations. Aside from the Distance model, the Size model performed remarkably well in many ROIs, especially in ventral cortex. In the data-model correlations, the Size model performed significantly above chance in EVC (p=5.79E-6), LOC (p=.000723), and pMTG (p=.00641), and marginally so in VMT (p=.0268) and SPOC (p=.0440). Similarly, in the MDS plots, smaller stimuli clustered separately from larger stimuli in EVC and VMT, and marginally in LOC (**Table 3**). Notably, in these ventral ROIs, stimulus grouping according to small vs. large size was orthogonal to grouping by near vs. far distance. This pattern suggests that size representations in these areas were not determined by retinal extent. If the representations were driven by retinal extent, we should expect to observe a linear organization in the MDS arrangements, with near large objects at one end, far small objects at the other, and the remaining stimuli distributed accordingly in between; however, this was not the case, even in strongly retinotopic areas of the cortex such as EVC. Rather, in EVC for instance, the position in the MDS plot of the near real hammer was roughly equidistant from the near real scissors as from the far real scissors, whereas if the representations were purely retinotopic, the near real hammer should be considerably closer to the near real scissors than the far real scissors, given the distance-driven difference in retinal extent between the two scissors. This orthogonal organization by distance and size was most evident in the ventral visual pathway and early visual cortex, where we might otherwise expect representations to be most strongly determined by retinal extent (Wandell and Winawer, 2011). Similarly, in the data-model correlations, the model RDMs for size and distance were orthogonal to each other. That both models performed significantly above chance in these regions also indicates that there are separable representations of object distance and size that cannot be explained fully by retinal extent.

Taken together, egocentric distance predominated as an organizing principle for cortical object representations across most, if not all, of the ROIs we tested. Yet object responses in many ROIs also carried information about other stimulus characteristics, including identity, grip type, and stimulus size.

### Egocentric distance modulates object representations in dorsal more than ventral cortex

The preceding analyses treat distance as one stimulus characteristic among many, yet the powerful overarching effect of the Distance model in the data-model correlations, together with the distinct clustering of stimulus representations in the near vs. far positions in the MDS plots, suggest that distance is a principal organizing factor of object representations. Yet it could be that representations of some object characteristics were highly distinctive across the near and far positions, but these differences were masked because the data were pooled across the near and far distances. For example, if representations of ‘hammers’ presented nearby are sufficiently different from those of ‘hammers’ presented at a distance, then an Identity model would not perform well in the aggregated data; however, it would be incorrect to conclude that stimulus identity is not represented at one or both locations. While an interaction model could hypothetically address the issue of distinct representations across the distance conditions, it is difficult to know which interaction model to test—and there are many such models that could be evaluated, even for a single characteristic, such as identity, let alone for multiple characteristics. Furthermore, there are clear *a priori* theoretical and empirical reasons for exploring object representations separately for stimuli at different egocentric locations. From a theoretical perspective, real objects presented at a near egocentric position are uniquely relevant for manual interaction, whereas this is less so for real objects presented at a far position (because they are outside of reach), and for pictures at either location (because pictures do not afford manual actions). From an empirical perspective there is increasing evidence that real objects elicit different responses from pictures specifically when the objects are actable (Bushong et al., 2010; Gomez et al., 2018; Fairchild et al., 2021; Snow and Culham, 2021; Snow et al., 2023). As such, a model for Display Format should be predicted to perform well only for stimuli presented at the near position because neither the real objects nor the pictures lend themselves to action when they are far away. For similar reasons, we might expect that object representations in dorsal regions are more affected by egocentric distance than those in ventral regions (Goodale and Milner, 1992). Therefore, we examined the effect of distance on representations of display format, identity, grip type, and size, separately for stimuli presented at near vs. far locations.

Figure 7 displays the mean Spearman’s Rank Correlation between the Display Format, Grip Type, Identity and Size models, and the Near (upper panel) and Far (lower panel) data RDMs. Model performance is shown separately for ROIs in ventral and dorsal cortex. Figure 8 displays the corresponding MDS plots for stimuli presented in the near and far locations. Inspection of Figure 7 shows that, overall, while many of the models performed significantly above chance for stimuli presented at the near location in both ventral and dorsal cortex, the same was not the case for stimuli presented at the far location, where model performance remained unaffected in ventral ROIs but failed to reached significance in any of the dorsal ROIs. A dependence on nearness in dorsal but not ventral ROIs was also evident in clustering by object features in the MDS plots.

**Figure 7:**
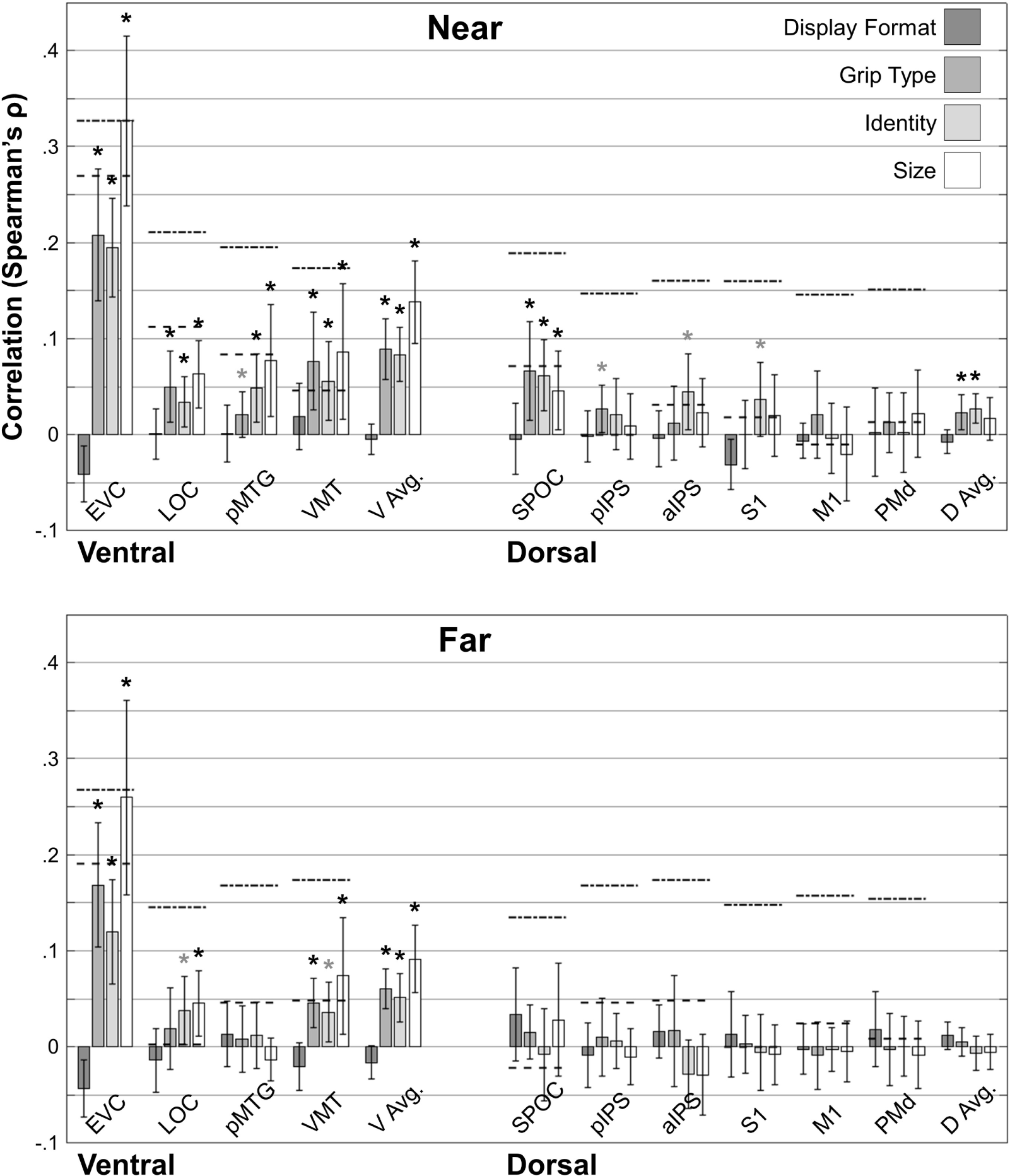
Data-model correlations as in Figure 5, calculated separately for the Near and Far conditions.

**Figure 8:**
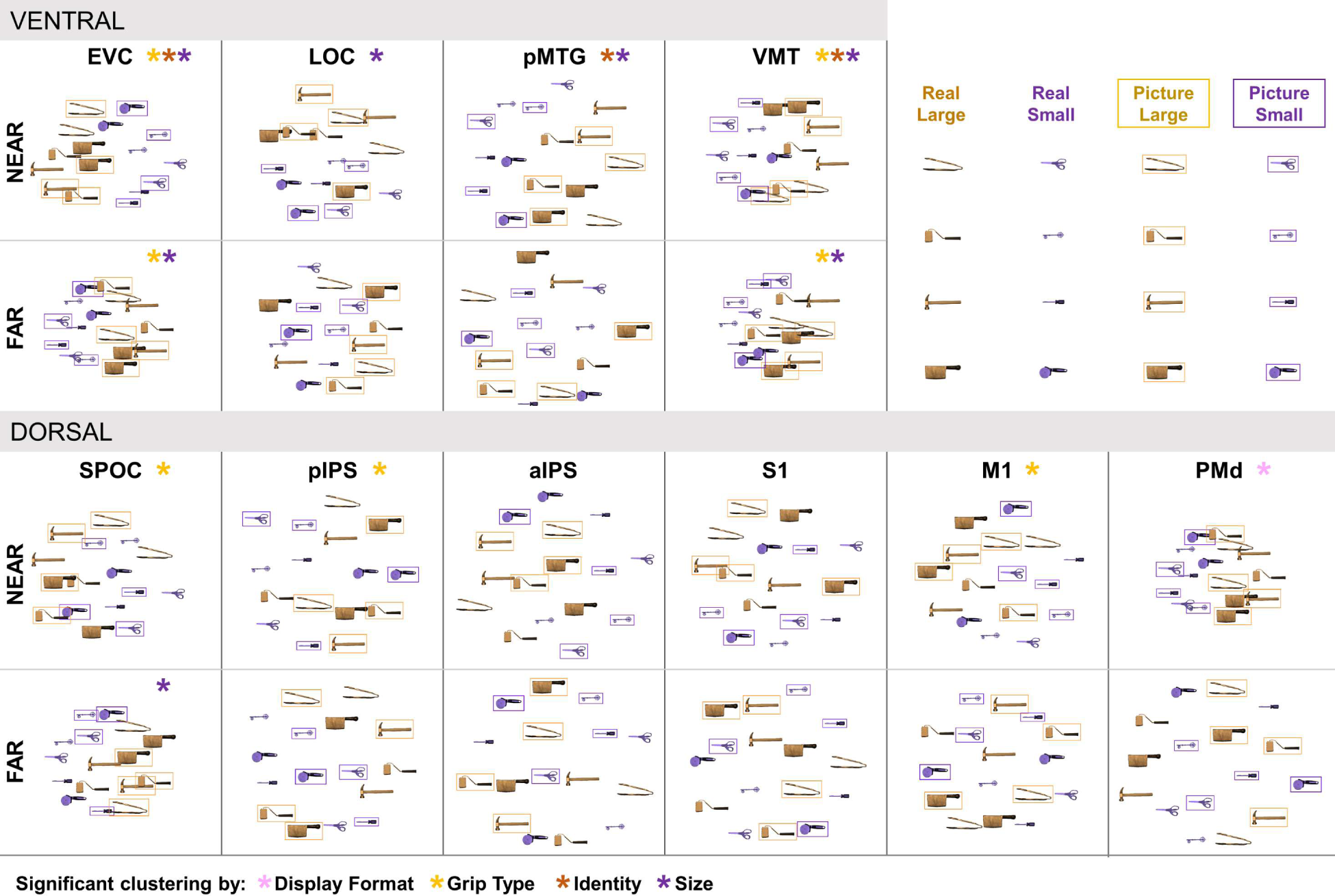
MDS plots as in Figure 6, created separately for the Near and Far conditions. Colors indicate stimulus size category. The four larger stimuli are shown in gold, while the four smaller stimuli are shown in purple. Stimuli presented as pictures are shown in rectangular frames, while stimuli presented as real objects are shown without such frames.

Looking first at the results for the Near condition in ventral ROIs, the data-model correlation analyses revealed that the models that performed well in all ROIs were Grip Type (EVC: p=2.14E-6, LOC: p=5.68E-3, pMTG: p=. 0396, VMT: p=2.72E-3), Identity (EVC: p=8.77E-8, LOC: p=6.81E-3, pMTG: p= 4.86E-3, VMT: p=5.02E-3), and Size (EVC: p=1.40E-7, LOC: p=7.02E-4, pMTG: p=6.11E-3, VMT: p=9.44E-3). Likewise, in the MDS plots, representations of near stimuli in ventral ROIs clustered by Grip Type in EVC and VMT, and by Identity and Size in all ROIs (**Table 4**). Conversely, the model for Display Format was not significant in any ventral ROI, nor did the representations evince any clustering by Display Format in the MDS plots.

**Table 4:** *P*-values for clusters in the MDS plots of Figure 8 for the Near condition.

| ROI | Display Format | Grip Type | Identity | Size |
| --- | --- | --- | --- | --- |
| EVC | 0.780 | .00105* | 7.58E-06* | 2.80E-13* |
| LOC | 0.780 | .332 | .0270* | 0.000970* |
| pMTG | 0.728 | .229 | .000921* | 1.22E-07* |
| VMT | 0.717 | .00523* | .00654* | 5.40E-07* |
| SPOC | 0.577 | 3.99E-05* | .0132* | 0.0827 |
| pIPS | 0.617 | .00196* | .594 | 0.472 |
| aIPS | 0.745 | .112 | .0501 | 0.0526 |
| S1 | 0.695 | .763 | .205 | 0.167 |
| M1 | 0.623 | .000827* | .126 | 0.0302* |
| PMd | 0.00134* | .0333* | .622 | 0.388 |

Next, for the Near condition in dorsal ROIs, performance of the Grip Type (SPOC: p=7.25E-3, pIPS: p=.0177), Identity (SPOC: p=1.17E-3, aIPS: p=.0139, S1: p=.0305), and Size (SPOC: p=.0151, pIPS: p=.0177) models was significant (or marginally so) in many ROIs, as was clustering according to these object features in the MDS plots (**Table 4**). Unlike in the ventral ROIs, yet matching the previous analysis of the undivided dataset, there was significant clustering by Display Format in the MDS plot for PMd.

Next, looking at the results for the Far condition in ventral cortex, the models for Grip Type (EVC: p=1.49E-5, VMT: p=6.67E-4), Identity (EVC: p=8.82-5, LOC: p=.0198, VMT: p=.0131), and Size (EVC: p=1.72E-5, LOC: p=5.98E-3, VMT: p=9.50E-3) continued to perform relatively well. Stimulus representations in the MDS plots continued to cluster by Grip Type and Size in EVC and VMT (**Table 5**). Overall, these data suggest that in ventral ROIs, stimulus distance has little influence on the representations of grip type, identity, and size (i.e., they show representational invariance, cf. Xu, 2018). However, as we observed in the Near condition, there was no apparent representation of Display Format in either the data-model correlations or the MDS arrangements.

**Table 5:** *P*-values for clusters in the MDS plots of Figure 8 for the Far condition.

| ROI | Display Format | Grip Type | Identity | Size |
| --- | --- | --- | --- | --- |
| EVC | 0.788 | .000910* | 0.0210* | 4.71E-12* |
| LOC | 0.305 | .612 | .744 | .353 |
| pMTG | 0.320 | .782 | .948 | .858 |
| VMT | 0.731 | .00610* | .240 | 2.41E-06* |
| SPOC | 0.223 | .155 | .226 | 6.69E-6* |
| pIPS | 0.822 | .699 | .284 | .204 |
| aIPS | 0.0445* | .0593 | .921 | .744 |
| S1 | 0.568 | .518 | .317 | .841 |
| M1 | 0.166 | .770 | .864 | .848 |
| PMd | 0.0391* | .599 | .852 | .844 |

Finally, the results for the Far condition in dorsal cortex revealed that none of the models performed above chance in any ROI, nor was there any significant clustering in the MDS plots for far stimuli by Grip Type, Identity, or Display Format in any ROI (**Table 5**). The only significant feature-based clustering for stimuli presented in the far position was for Size in SPOC.

Taken together, these data suggest that positioning stimuli out of reach attenuates the representation of object characteristics in dorsal ROIs, consistent with the dorsal pathway’s putative role in representing egocentric space around the body (Mishkin and Ungerleider, 1982; Mishkin et al., 1983; Goodale and Milner, 1992; Vallar et al., 1999; Galati et al., 2000). This contrasts with the results for ventral ROIs, where egocentric distance did not consistently modulate representation of stimulus features.

### Size representations depend on a combination of distance and display format in dorsal but not ventral cortex

In the analyses reported above, we found that stimulus size was consistently represented in multivariate patterns in ventral ROIs, both in our analysis of the undivided dataset, and in our separate analyses of the Near and Far conditions. Size was represented in ventral ROIs for stimuli presented at either distance. Conversely, we found that stimulus size was rarely, if at all, represented in the multivariate patterns in dorsal ROIs. This was the case both for stimuli presented in the near and in the far position. Given the dorsal pathway’s putative role in perception in the service of action (Erlikhman et al., 2018; Freud et al., 2020), we might expect size representation in these regions only for stimuli that offer the opportunity for action. If this is the case, then size representations in dorsal ROIs in our experiment should be apparent when the stimuli are real objects in the near position, but not when they are real objects in the far position, or pictures at either position. Because previous neuroimaging studies have found that object responses in the ventral pathway remain invariant across changes in the size or format in which a stimulus is displayed (Kourtzi and Kanwisher, 2000; Grill-Spector et al., 2001; Vuilleumier et al., 2002), size representations in ventral ROIs in our experiment should be similar irrespective of display format or distance, in contrast with our prediction for dorsal cortex.

To test these predictions, we examined size clustering in the MDS patterns separately for the Real Near, Real Far, Picture Near, and Picture Far conditions. In ventral ROIs, we found significant clustering by stimulus size, a pattern relatively consistent across both display formats and both egocentric positions (**Table 6**). In certain dorsal ROIs, clustering by stimulus size only emerged when stimuli were presented as real objects in the near position; however, this was only apparent in two of the most anterior ROIs, S1 and PMd. To illustrate this pattern, we show the MDS plots for representative ROIs in Figure 9 (MDS plots for the remaining ROIs available in **Supplementary Figure S6**). For example, the four largest and the four smallest stimuli are linearly separable for EVC in all conditions, whereas this is the case in S1 only for the Real Near condition.

**Table 6:** *P*-values for clusters in the MDS plots of Figure 9.

| ROI | Real Near | Real Far | Picture Near | Picture Far |
| --- | --- | --- | --- | --- |
| EVC | .00100* | .00273* | .000571* | .00111* |
| LOC | .0215* | .299 | .0319* | .307 |
| pMTG | .0291* | .541 | .00500* | .191 |
| VMT | .0103* | .00297* | .0386* | .0235* |
| SPOC | .509 | .0216* | .801 | .0175* |
| pIPS | .875 | .00662* | .830 | .607 |
| aIPS | .391 | .548 | .477 | .558 |
| S1 | .00789* | .883 | .390 | .546 |
| M1 | .682 | .345 | .0634 | .859 |
| PMd | .00889* | .505 | .572 | .895 |

**Figure 9:**
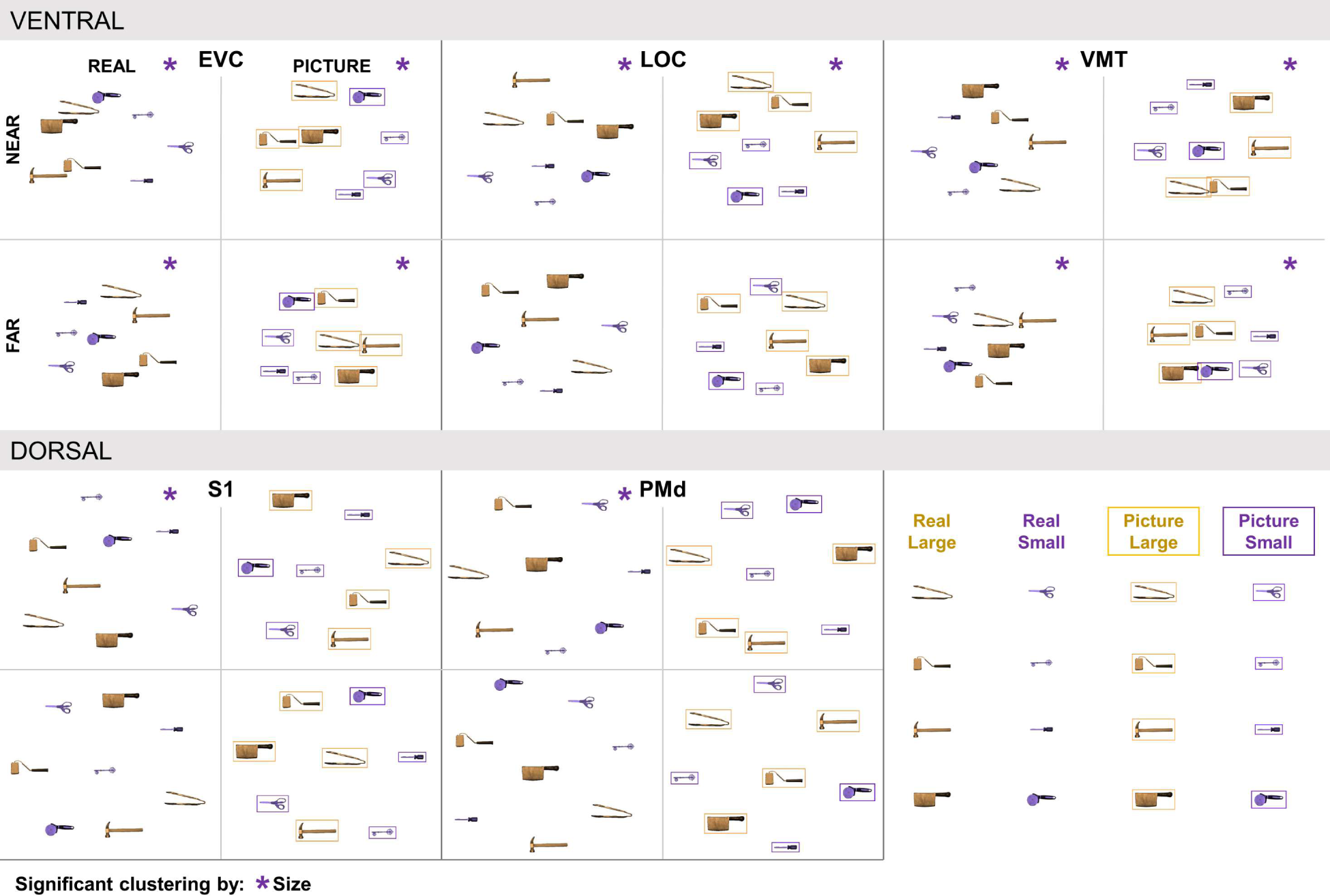
MDS plots created separately for the Real Near, Real Far, Picture Near, and Picture Far conditions. Colors indicate stimulus size category. The four larger stimuli are shown in gold, while the four smaller stimuli are shown in purple. For each ROI, the Real Near plot is displayed in the upper left, Real Far in the lower left, Picture Near in the upper right, and Picture Far in the lower right.

## Discussion

Here, we used fMRI to examine how display format and egocentric distance influenced object responses in human cortex. Participants directly viewed real objects and matched pictures that were positioned either nearer to the observer, or farther from the observer.

We found that egocentric distance was both a widespread determinant of univariate BOLD amplitudes, and a principal organizing factor of multivariate response patterns; notably, the effects of distance were typically apparent across both dorsal and ventral cortex. Analysis of response amplitudes revealed that many cortical regions responded more strongly to near than far stimuli, consistent with previous work showing that egocentric distance modulates BOLD amplitudes (Gallivan et al., 2009, 2011; Cavina-Pratesi et al., 2010). Our multivariate analyses extend these findings by showing that egocentric distance also modulates object response patterns. Stimulus distance modulated response patterns not only in dorsal cortex, but also in ventral cortex. For example, in many dorsal regions, object characteristics such as display format, size, identity, and grip type were apparent in the representations for near but not far stimuli. Although the egocentric position of stimuli in the ‘near’ and ‘far’ positions in our study differed by only 38 cm, the ‘near’ stimuli fell within participant’s reach, while those in the ‘far’ position lay beyond reach. Whether our findings reflect sensitivity to distance per se (i.e., whereby object responses should change gradually with increasing distance), or sensitivity to object reachability (i.e., where object responses should exhibit a sharp discontinuity at the boundary of peripersonal space) could be addressed in future work by using parametric distance manipulations. We observed striking effects of distance on responses to both real objects *and* size-matched pictures of those objects—a result that seems surprising given that pictures convey ambiguous distance information and are not inherently graspable. Yet it is possible that distance influenced cortical responses to objects depicted in pictures because printed pictures have an unusual dual nature (DeLoache et al., 1998): They are objects in themselves (which *are* graspable), and at the same time they are representations of something else. The objects depicted in our pictures were also matched in size to their real-world counterparts. In these two respects, our stimuli contrast markedly with those used in previous fMRI studies, where the projected images convey little, if any, information to the observer about the distance, location, or real-world size of the depicted object, and where size is further obscured through stimulus scaling. In many fMRI studies, objects that are typically very small in the real world are presented with the same retinal extent as objects that are orders of magnitude larger in the real world. Future studies could compare responses between digital images, printed pictures, and real objects to determine the extent to which stimulus physicality influences format-related effects.

As with egocentric distance, object size was widely represented across ventral and dorsal cortex. In many brain regions, size was represented orthogonally to distance in the response patterns, suggesting that size and distance are represented independently in human cortex, and that the representations cannot be explained by differences in retinal extent. These findings are consistent with electrophysiological evidence that stimulus size and distance are reflected in cortical responses to real objects even when their retinal extent is held constant (Noviello et al., 2024). Our results align with previous studies showing topographic organization in ventral cortex by the expected real-world size of objects (Konkle and Oliva, 2012; Konkle and Caramazza, 2013). Yet unlike in previous studies using projected images, for which a given retinal extent is compatible with a range of size-distance combinations, our physical stimuli have definite sizes and egocentric distances that are not inferred based on prior expectations. Interestingly, size was represented for nearby real objects in somatomotor cortex even though participants did not manually interact with the stimuli. Indeed, some of the regions that represented stimulus size in ventral and dorsal cortex have also been implicated in representing object weight (Chouinard et al., 2005; Gallivan et al., 2014; Schwettmann et al., 2019) and length (Fabbri et al., 2016). Because size, weight, and elongation are typically correlated, the “size” representations we observed could reflect any of these characteristics; future studies could tease apart these alternatives by using larger stimulus sets that allow different physical features to be decorrelated. Furthermore, given that real objects (but not pictures) are processed according to their weight (Holler et al., 2020), future studies could examine whether, and where, weight emerges in the associated neural representations (Chouinard et al., 2009; Fairchild & Snow, 2020).

The format in which objects were displayed had a greater influence on fMRI amplitudes than on representational patterns. Amplitude responses were higher for real objects than for pictures in lateral occipital and parietal cortices, dovetailing with previous fMRI studies (Snow et al., 2011; Freud et al., 2018). Critically, we advance on previous findings by showing that format-related differences in LOC are not limited to repetition suppression (Snow et al., 2011) but also extend to response amplitudes, and that format-related differences in parietal cortex are not limited to action (Freud et al., 2018) but are also observed during perception. Importantly, we found that response amplitudes to real objects were modulated by stimulus distance more than responses to pictures in much of ventral and dorsal cortex (Gomez et al., 2018; Snow et al., 2023). Display format did not influence response patterns as widely as response amplitudes, although display format was consistently represented in dorsal premotor cortex, particularly for nearby stimuli. Taken together, these findings suggest that although certain brain regions are driven more strongly by real objects than by pictures, most of human cortex does not have distinct representations for pictorial stimuli –perhaps because pictures emerged relatively recently in hominid evolution (Aubert et al., 2019) in contrast to real objects, for which three-dimensionality, actability, weight, and other physical features have had consequences for behavior and survival for millions of years (Cisek and Kalaska, 2010).

Finally, we found that stimulus distance affected object representations differently in ventral compared to dorsal cortex. In ventral cortex, representations of identity, size, and grip type were largely unaffected by stimulus distance, dovetailing with previous findings that ventral representations remain stable despite changes in stimulus size and format (Kourtzi and Kanwisher, 2000; Grill-Spector et al., 2001; Vuilleumier et al., 2002), and across contextual changes (Vaziri-Pashkam and Xu, 2017, 2019; Xu, 2018). In dorsal cortex, however, identity, size, and grip type were represented when stimuli were nearby, but not when they were farther away. Furthermore, two somatomotor dorsal ROIs, S1 and PMd, represented stimulus size only for real objects within reach, scaffolding findings from previous behavioral (Bushong et al., 2010; Gomez et al., 2018; Snow et al., 2023) and EEG (Fairchild et al., 2021) studies showing that reachability modulates responses to real objects, but not to pictures. When a stimulus cannot be acted upon, either because it is out of reach or is merely a picture, representation of stimulus size and weight may not serve somatomotor determination of grip apertures (Holmes and Heath, 2013; Ozana et al., 2018; Ozana and Ganel, 2019) and grip force (Chouinard et al., 2005; Loh et al., 2010; van Nuenen et al., 2012). These results, along with previous work showing that real objects are more likely than pictures to be processed according to their size and weight (Holler et al., 2020), and that the size of real objects is a more important cue to object identity than for pictures (Holler et al., 2019; Sensoy et al., 2021), support recent proposals that dorsal cortex is uniquely involved in perception of physical object features relevant for action (Erlikhman et al., 2018; Freud et al., 2020). Our results build upon this view by suggesting not only that dorsal cortex contributes to perception, but also that ventral cortex contributes to spatial cognition. In ventral cortex, while distance had little effect on individual stimulus characteristics such as size, activation patterns differed substantially for near versus far stimuli. Our results parallel findings from studies of image perception which have shown that distance modulates responses in ventral cortex when images of objects (Amit et al., 2012) or scenes (Persichetti and Dilks, 2016) include salient pictorial distance cues. Representation of distance could facilitate the ventral pathway’s role in object recognition, as different objects are more likely to be encountered at different distances (Wang et al., 2024). Taken together, these results support recent frameworks that emphasize the functional integration between the dorsal and ventral visual pathways (Freud et al., 2017, 2020; Almeida et al., 2018; Wurm and Caramazza, 2022).

Our findings underscore that the stimulus characteristics represented in brain activity depend critically on the information conveyed by the stimuli themselves. Stimuli that convey unambiguous cues about physical properties, such as distance and size, are more likely to reveal representations of these stimulus characteristics across the cortex. Experimental stimuli that convey rich, multidimensional object characteristics are likely to engage the brain in a way that most closely resembles naturalistic vision.

## Acknowledgments

This research was supported by grants from the National Eye Institute of the National Institutes of Health (NIH, grant number R01EY026701 to J. C. S.) and the National Institute of General Medical Sciences of the NIH (grant numbers P20GM103650 and P30GM145646). We thank Marlene Behrmann and Michael Rudd for their feedback on statistical analyses, Kallie McDonald for assistance with supplementary figure generation, Carissa Romero for assistance with data collection, and Larry Messier for technical neuroimaging assistance.

## Supplementary Figures

**Figure S1:**
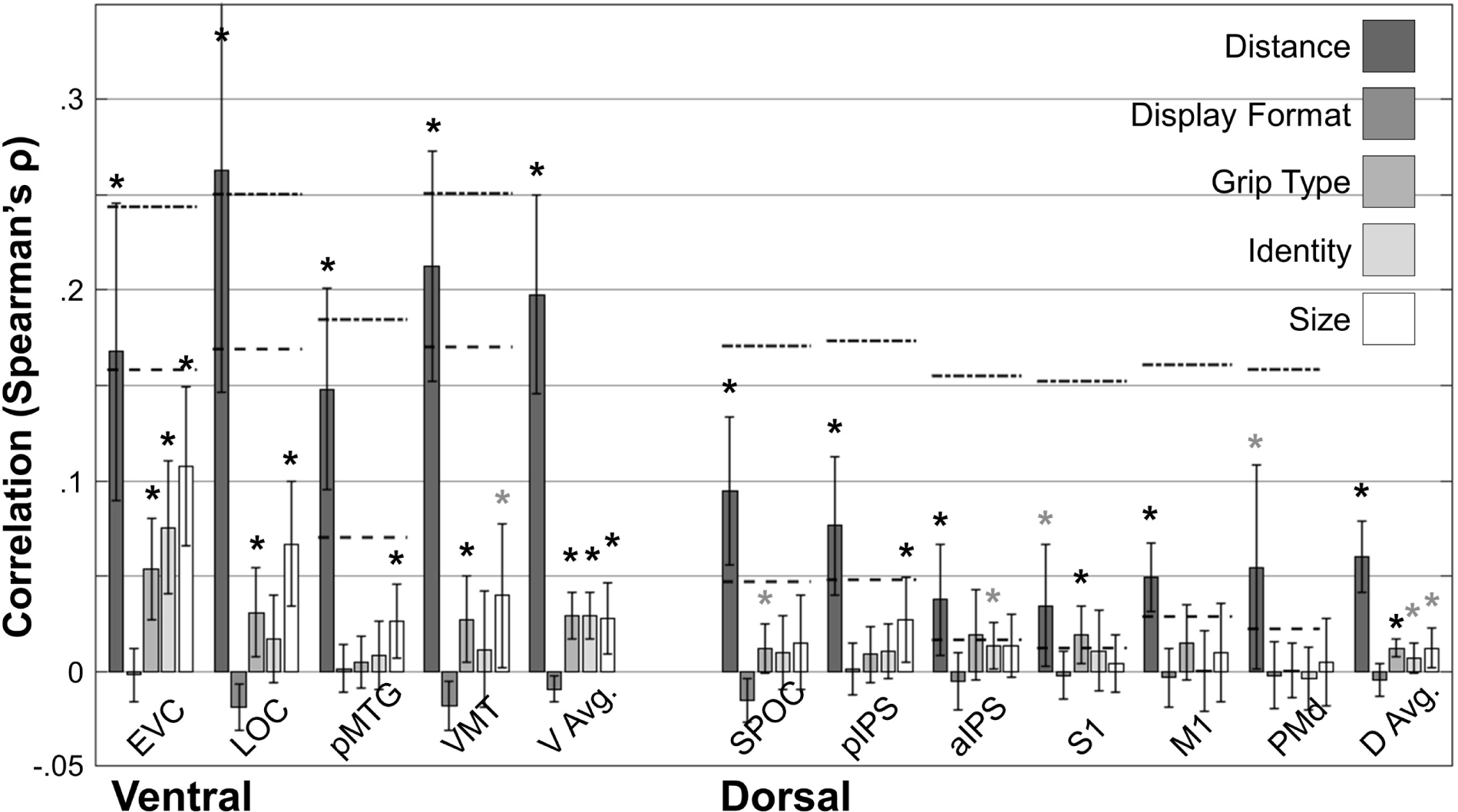
Right-hemisphere data-model correlations for the undivided dataset.

**Figure S2:**
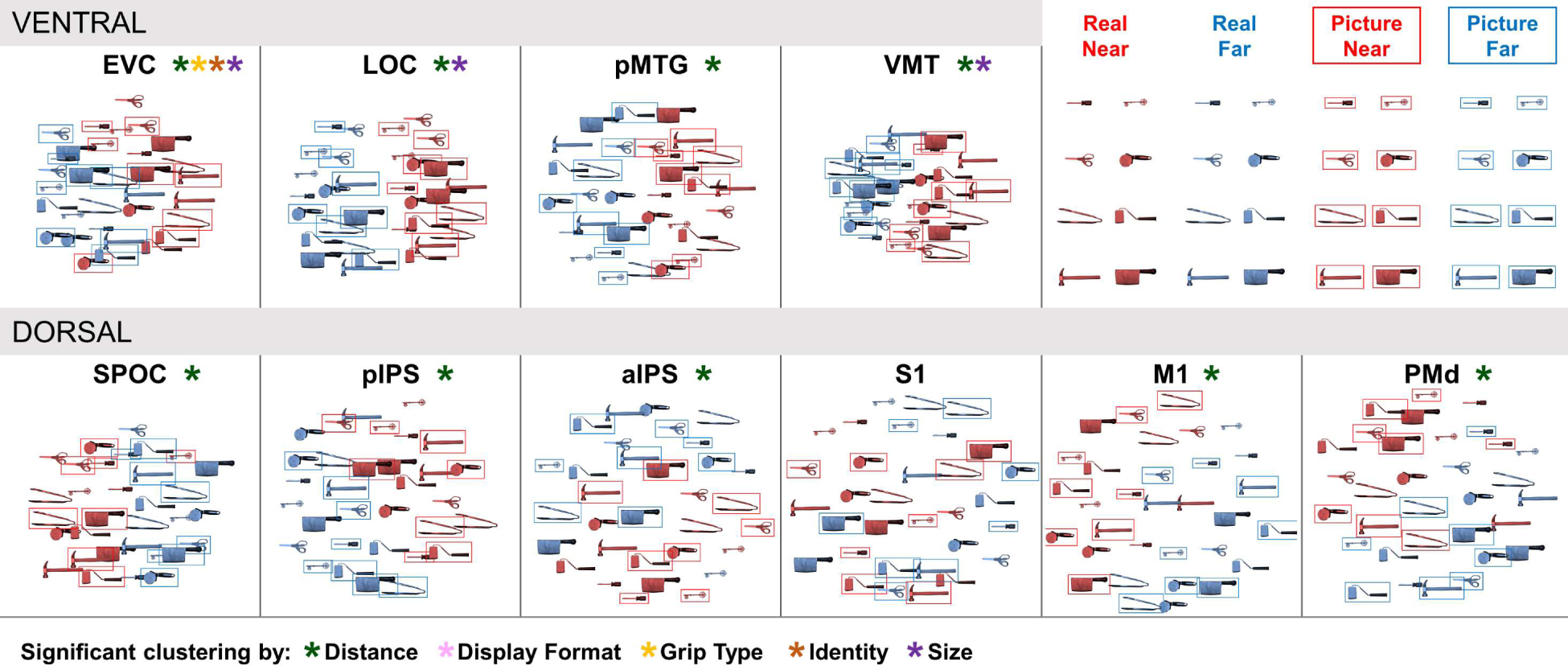
Right-hemisphere MDS plots for the undivided dataset.

**Figure S3:**
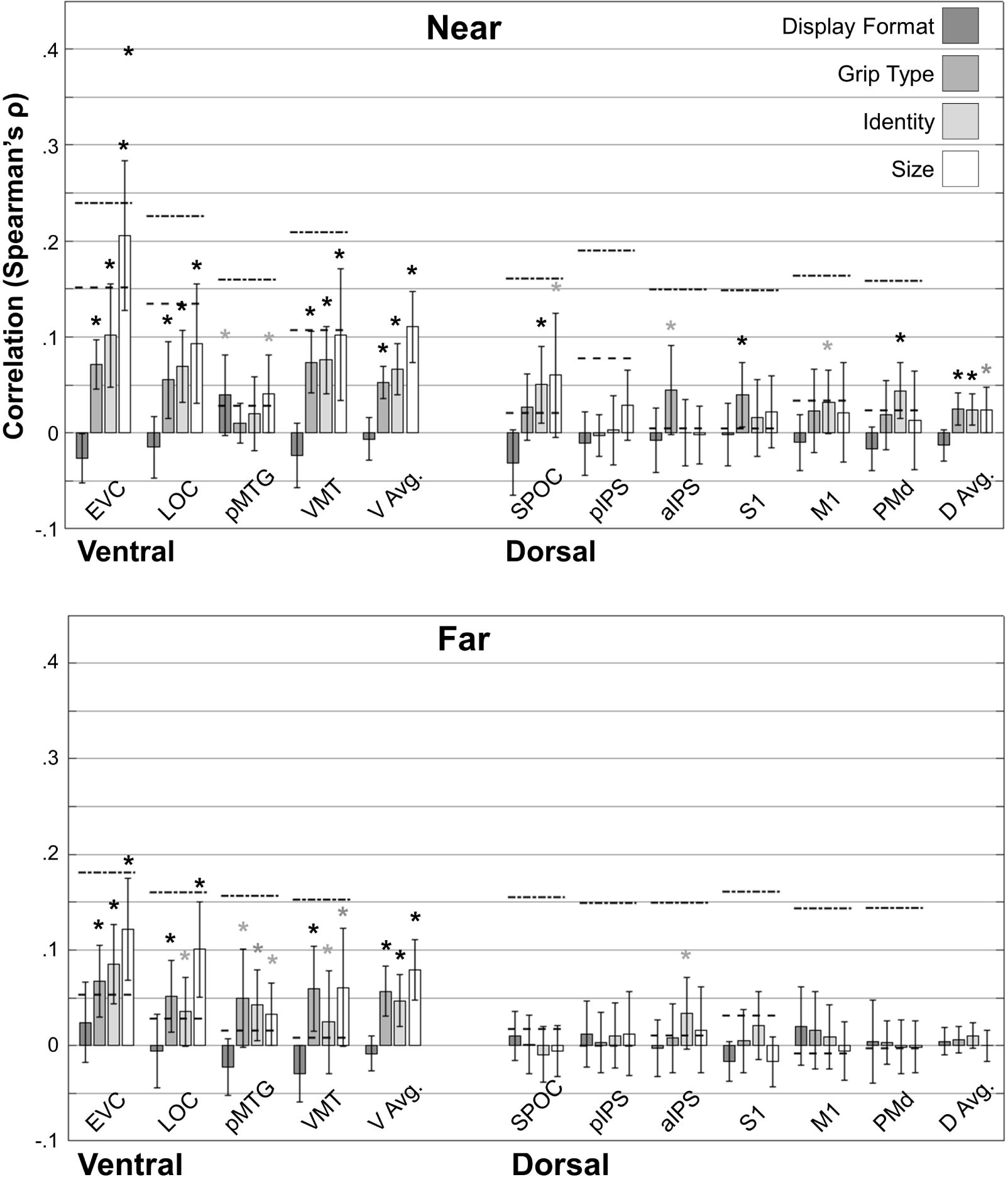
Right-hemisphere data-model correlations calculated separately for the Near and Far conditions.

**Figure S4:**
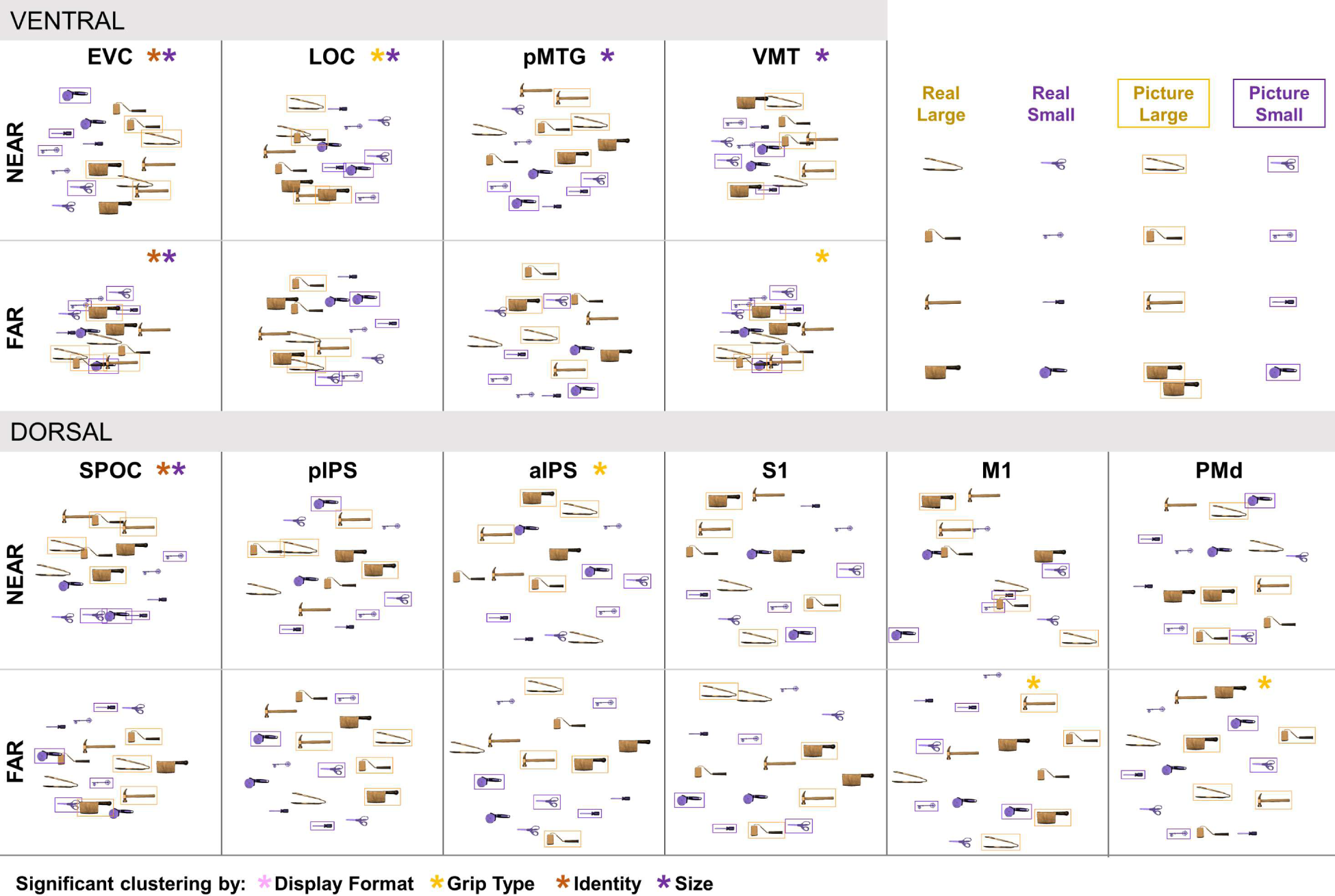
Right hemisphere MDS plots created separately for the Near and Far conditions.

**Figure S5:**
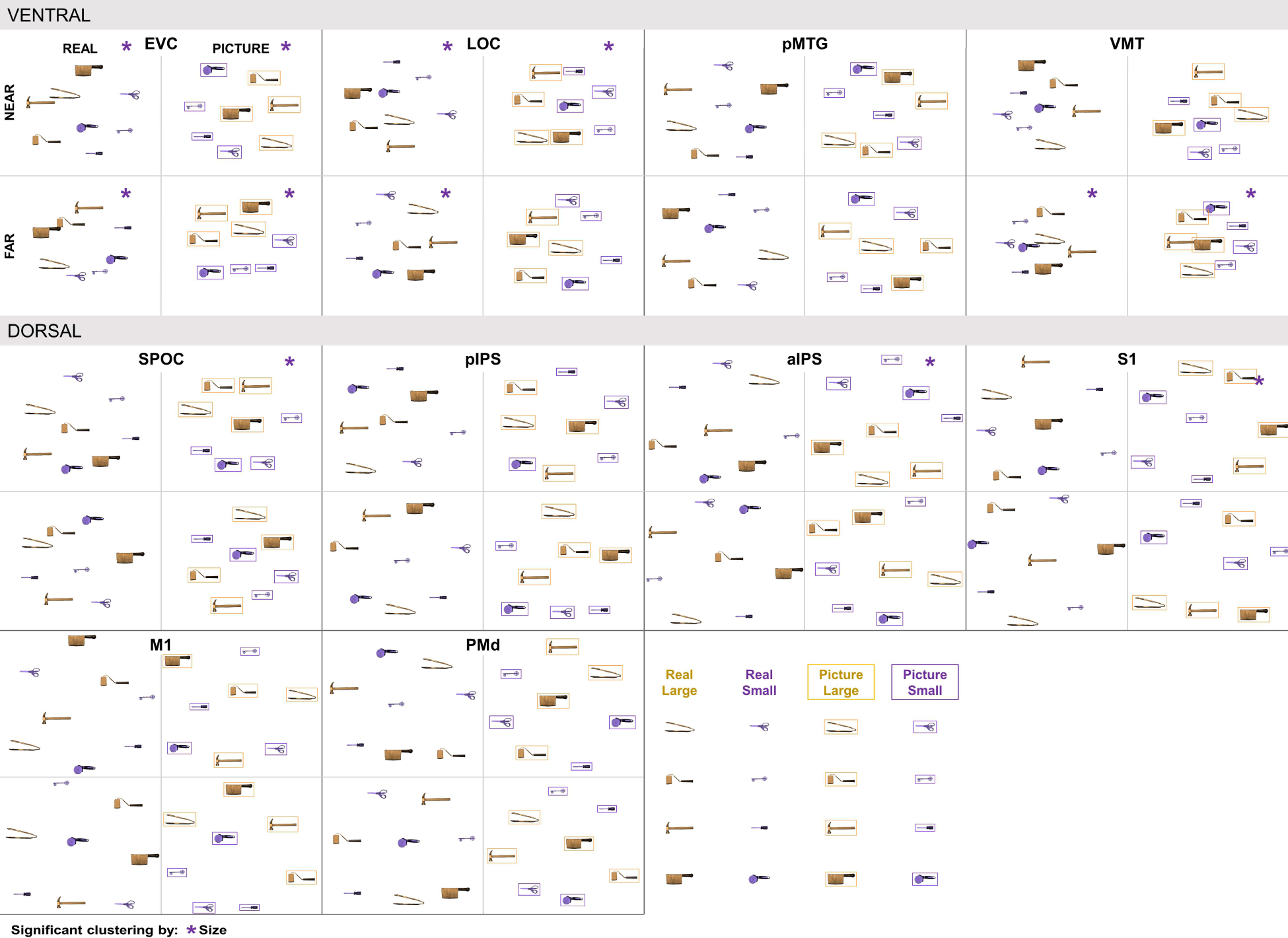
Right-hemisphere MDS plots created separately for the Real Near, Real Far, Picture Near, and Picture Far conditions.

**Figure S6:**
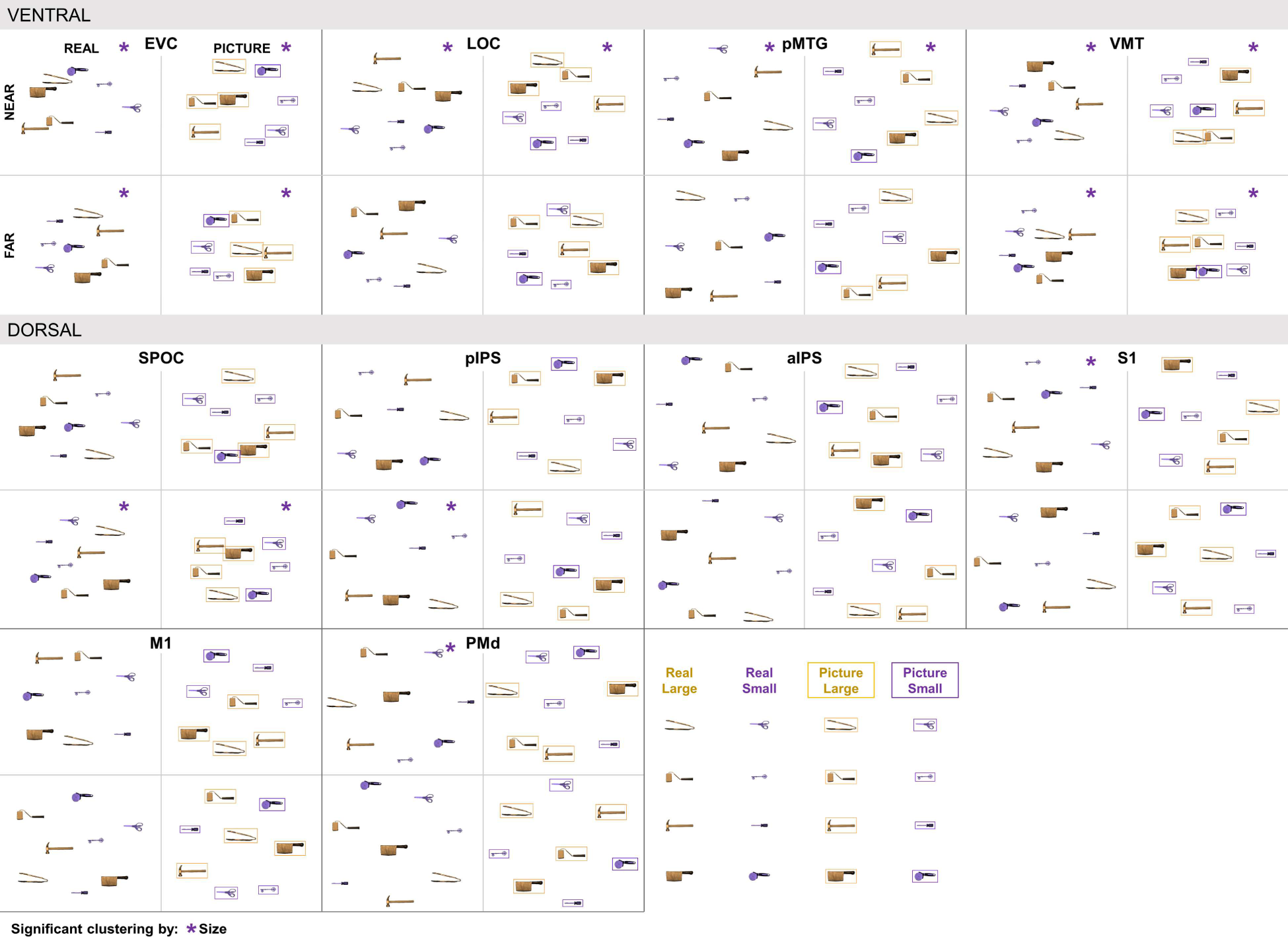
Left-hemisphere MDS plots created separately for the Real Near, Real Far, Picture Near, and Picture Far conditions. ROIs not shown in Figure 9 are shown here.

## Supplementary Tables

**Table S1.**
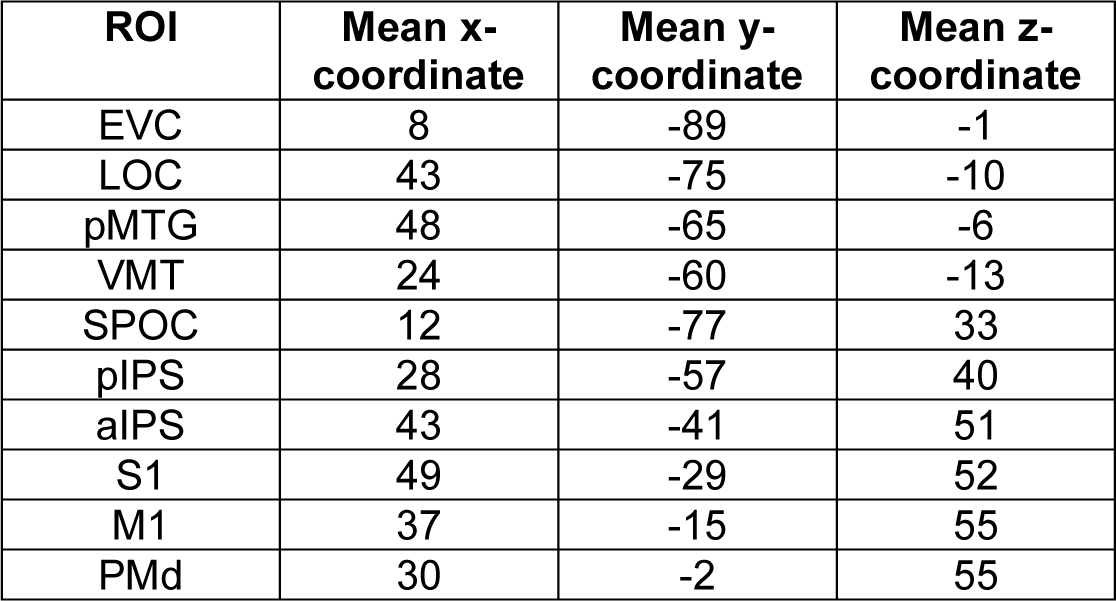
MNI coordinates for right-hemisphere regions of interest used in multivariate analyses. Although aIPS may not be expected to have a functional analogue in the right hemisphere, we nonetheless defined an analogous right-hemisphere region using similar anatomical criteria.

**Table S2:** *P*-values for clusters in the MDS plots of Figure S2. *P*-values below 1E-15 are reported as 0.

| ROI | Distance | Display Format | Grip Type | Identity | Size |
| --- | --- | --- | --- | --- | --- |
| EVC | 0* | 0.837 | 0.000699* | 0.00114* | 4.35E-05* |
| LOC | 0* | 0.762 | 0.0227* | 0.109 | 0* |
| PMTG | 0* | 0.730 | 0.544 | 0.680 | 0.107 |
| VMT | 0* | 0.813 | 0.125 | 0.591 | 0.0061* |
| SPOC | 0* | 0.730 | 0.254 | 0.838 | 0.297 |
| pIPS | 0* | 0.429 | 0.773 | 0.647 | 0.217 |
| aIPS | 0* | 0.838 | 0.0271* | 0.445 | 0.210 |
| S1 | 0.147 | 0.627 | 0.0588 | 0.323 | 0.539 |
| M1 | 0* | 0.826 | 0.151 | 0.580 | 0.799 |
| PMd | 0* | 0.431 | 0.469 | 0.516 | 0.562 |

**Table S3:** *P*-values for clusters in the MDS plots of Figure S4 for the Near condition.

| ROI | Display Format | Grip Type | Identity | Size |
| --- | --- | --- | --- | --- |
| EVC | 0.820 | 0.124 | 5.30E-05* | 1.45E-10* |
| LOC | 0.158 | 0.00665* | 0.0552 | 3.14E-07* |
| PMTG | 0.281 | 0.753 | 0.119 | 0.009912* |
| VMT | 0.815 | 0.195 | 0.0790 | 0.000578* |
| SPOC | 0.740 | 0.200 | 0.000506* | 2.08E-06* |
| pIPS | 0.774 | 0.624 | 0.545 | 0.679 |
| aiPS | 0.678 | 0.00344* | 0.175 | 0.211 |
| S1 | 0.352 | 0.301 | 0.823 | 0.425 |
| M1 | 0.787 | 0.299 | 0.156 | 0.692 |
| PMd | 0.751 | 0.801 | 0.0930 | 0.754 |

**Table S4:** *P*-values for clusters in the MDS plots of Figure S4 for the Far condition.

| ROI | Display Format | Grip Type | Identity | Size |
| --- | --- | --- | --- | --- |
| EVC | 0.777 | 0.148 | 0.0120* | 0.000185* |
| LOC | 0.305 | 0.612 | 0.554 | 0.744 |
| PMTG | 0.320 | 0.782 | 0.745 | 0.948 |
| VMT | 0.731 | 0.00610* | 0.112 | 0.240 |
| SPOC | 0.705 | 0.434 | 0.915 | 0.370 |
| pIPS | 0.344 | 0.289 | 0.206 | 0.105 |
| aiPS | 0.754 | 0.814 | 0.0131* | 0.564 |
| S1 | 0.451 | 0.121 | 0.0398* | 0.729 |
| M1 | 0.727 | 0.00193* | 0.168 | 0.131 |
| PMd | 0.659 | 0.000224* | 0.0730 | 0.196 |

**Table S5:** P-values for clusters in the MDS plots of Figure S5.

| ROI | Real Near | Real Far | Picture Near | Picture Far |
| --- | --- | --- | --- | --- |
| EVC | 0.00136* | 0.00139* | 0.00566* | 0.0110* |
| LOC | 0.0177* | 0.00482* | 0.0319* | 0.307 |
| PMTG | 0.514 | 0.374 | 0.748 | 0.754 |
| VMT | 0.0937 | 0.0148* | 0.344 | 0.0118* |
| SPOC | 0.416 | 0.862 | 0.0301* | 0.786 |
| pIPS | 0.642 | 0.403 | 0.646 | 0.147 |
| aiPS | 0.876 | 0.875 | 0.00945* | 0.630 |
| S1 | 0.536 | 0.876 | 0.0228* | 0.468 |
| M1 | 0.576 | 0.876 | 0.878 | 0.0710 |
| PMd | 0.874 | 0.870 | 0.678 | 0.826 |

